# LncRNA analyses reveal increased levels of non-coding centromeric transcripts in hepatocellular carcinoma

**DOI:** 10.1101/2021.03.03.433778

**Authors:** Anamaria Necsulea, Philippe Veber, Tuyana Boldanova, Charlotte K Y Ng, Stefan Wieland, Markus H Heim

## Abstract

The search for new biomarkers and drug targets for hepatocellular carcinoma (HCC) has spurred an interest in long non-coding RNAs (lncRNAs), often proposed as oncogenes or tumor suppressors. Furthermore, lncRNA expression patterns can bring insights into the global de-regulation of cellular machineries in tumors. Here, we examine lncRNAs in a large HCC cohort, comprising RNA-seq data from paired tumor and adjacent tissue biopsies from 114 patients. We find that numerous lncRNAs are differentially expressed between tumors and adjacent tissues and between tumor progression stages. Although we find strong differential expression for most lncRNAs previously associated with HCC, the expression patterns of several prominent HCC-associated lncRNAs disagree with their previously proposed roles. We examine the genomic characteristics of HCC-expressed lncRNAs and reveal an enrichment for repetitive elements among the lncRNAs with the strongest expression increases in advanced-stage tumors. This enrichment is particularly striking for lncRNAs that overlap with satellite repeats, a major component of centromeres. Consistently, we find increased non-coding RNA transcription from centromeres in tumors, in the majority of patients, suggesting that aberrant centromere activation takes place in HCC.

## Introduction

Following the realization that the human genome harbors thousands of non-coding RNA genes (Carninci *et al*, 2005), many of which have important cellular functions (Mattick & Makunin, 2006), a great deal of effort has been put into investigating the contributions of non-coding RNAs to cancer biology (Gutschner & Diederichs, 2012). In particular, the roles of long non-coding RNAs (lncRNAs) in cancer have been frequently scrutinized in the past decade. This category of non-coding RNAs (defined simply as RNA molecules that lack protein-coding capacity, at least 200 nucleotides long) comprises many transcripts with proven functions in gene expression regulation, genome stability or nuclear architecture (Engreitz *et al*, 2016). Numerous recent studies showed that lncRNA loci are part of the alterations that occur in cancer cells (Yan *et al*, 2015). Thus, studying lncRNAs is perceived as a promising path towards understanding the molecular mechanisms that underlie cancer onset. Ultimately, lncRNAs may prove to be valuable in the diagnosis process, or serve as therapeutic targets.

The search for novel disease biomarkers and drug targets, including lncRNAs, is understandably intensive for cancer types for which effective therapies are still lacking. This is the case for hepatocellular carcinoma (HCC), which is a major cause of cancer-related mortality world-wide (Yang & Roberts, 2010). As HCC is generally detected at late stages of tumor progression, surgical treatment options are unavailable for the majority of patients (Hartke *et al*, 2017). Several systemic therapies now exist, but they increase median patient survival by less than 1 year (Finn *et al*, 2020). Thus, developing new treatments for HCC is still an urgent need. With this aim, there has been extensive research aiming to identify the genic and non-genic functional elements that are altered in HCC compared to the healthy liver. Large-scale transcriptomics studies, comparing HCC samples with adjacent non-tumor tissue or with normal liver samples, identified hundreds of differentially regulated protein-coding genes and lncRNAs (Cui *et al*, 2017; Yang *et al*, 2017; Li *et al*, 2019; Jin *et al*, 2019; Unfried *et al*, 2019). Some of the lncRNAs associated with HCC through genome-wide comparative analyses were subject to further experimental investigations, aiming to elucidate their mechanisms of action and the consequences of their differential regulation in tumors. For some lncRNAs, there are now well-supported models for their behavior in HCC. This is the case for example for *HOTTIP*, a lncRNA that is strongly up-regulated in HCC, and which likely acts to enhance the expression of the neighboring genes by recruiting transcriptional co-activators (Quagliata *et al*, 2014; Pradeepa *et al*, 2017; Quagliata *et al*, 2018). However, for other lncRNAs experimental studies gave rise to conflicting results. For example, the *H19* lncRNA (known as a parentally imprinted regulator of placenta growth (Keniry *et al*, 2012)) was alternatively proposed to act as a tumor suppressor (Hao *et al*, 1993; Yoshimizu *et al*, 2008; Schultheiss *et al*, 2017) or as an oncogene (Matouk *et al*, 2007; Zhou *et al*, 2019) in various cancer types including HCC (Tietze & Kessler, 2020). Likewise, *MALAT1*, initially described as an abundant lncRNA associated with the presence of metastases (Ji *et al*, 2003), was first thought to promote tumor growth and invasion in breast cancer (Arun *et al*, 2016), but is now believed to be a tumor suppressor (Kim *et al*, 2018, 1). In HCC, *MALAT1* was mainly proposed to act as an oncogene (Hou *et al*, 2017, 1; Liu *et al*, 2019, 1; Chen *et al*, 2020), but there is no consensus on its mechanisms of action. This is also the case for most of the lncRNAs that have been associated with HCC, although experimental data is accumulating (Lanzafame *et al*, 2018). Thus, overall, the functions of lncRNAs in HCC and other cancers are still poorly understood.

Although we are still far from developing therapies that target lncRNAs in HCC, in the more immediate future, lncRNAs may prove to be useful as disease biomarkers, to help diagnose HCC at an earlier stage and to better classify molecular subtypes of tumors. For this purpose, large-scale transcriptomics comparisons that can identify differentially regulated lncRNAs in tumor tissues are a valid approach, even in the absence of additional functional experiments. Although such studies are abundant in the lncRNA literature, they are often restricted to small cohorts of patients, thus potentially failing to reproduce the full extent of the molecular heterogeneity of HCC (Boyault *et al*, 2007; Hoshida *et al*, 2009). Further work is still needed to understand what part lncRNAs play in the molecular characteristics of HCC.

Studying lncRNA expression patterns in HCC and other cancers is also a means to better understand the de-regulation of essential cellular machineries in tumors. Although many lncRNAs have important biological roles (Engreitz *et al*, 2016), there is strong evidence that, out of the tens of thousands of lncRNAs that are detected with sensitive transcriptome sequencing approaches in human tissues (Pertea *et al*, 2018; Iyer *et al*, 2015), most may be non-functional. This is indicated by their weak levels of evolutionary conservation (Necsulea *et al*, 2014; Washietl *et al*, 2014) and by their expression patterns, which are often restricted to tissues with open chromatin, permissive to spurious transcription (Soumillon *et al*, 2013; Darbellay & Necsulea, 2020). Other evidence supporting non-functionality, or even a deleterious effect of lncRNA transcription comes from their typical processing by the cellular machinery. LncRNA transcripts are generally inefficiently spliced and poly-adenylated, and are rapidly degraded by the RNA exosome (Melé *et al*, 2017; Schlackow *et al*, 2017). For certain classes of lncRNAs, transcription is normally tightly repressed by chromatin-modifying factors, and their de-repression leads to DNA replication stress and subsequently to cellular senescence, due to an overlap with DNA replication origins (Nojima *et al*, 2018). It is not clear yet to what extent similar principles apply to HCC and other cancers. However, the presence of high lncRNA levels in cancer cells may be a sign of a global de-regulation of the molecular machineries that normally keep deleterious transcription in check, even if individual lncRNAs are not “oncogenes” *sensu stricto*. This further highlights the need for detailed investigations of the patterns of lncRNA expression in cancer.

In this study, we set out to explore the patterns of lncRNA transcription in a large HCC cohort, comprising paired tumor and adjacent tissue biopsies from 114 patients. Our work stands out from previous efforts to characterize lncRNAs in HCC in several important ways. First, we take advantage of an extensive transcriptome resource, which covers a wide range of tumor progression stages and underlying liver diseases, and thus can provide a comprehensive overview of transcriptional de-regulation during HCC development. Importantly, our transcriptome dataset is derived from biopsies rather than tumor resections, and is thus likely more faithful to the *in vivo* physiological status of the tumors. Second, we perform a meta-analysis of the current literature on lncRNAs and HCC and we use our transcriptome collection to critically re-evaluate previous claims regarding lncRNA expression patterns in HCC. We can thus highlight the poor reproducibility of some prominent lncRNA-HCC associations. Third, rather than attempting to propose new candidate oncogene or tumor suppressor lncRNAs, we perform a detailed analysis of the genomic characteristics of de-regulated lncRNAs. We thus reveal an increase in repetitive-element derived lncRNA expression in tumor samples. In particular, we uncover a striking up-regulation of non-coding RNAs derived from centromeric satellite repeats. We discuss the functional implications of this apparent activation of centromeric chromatin in HCC tumors.

## Results

### Transcriptome dataset

We analyzed the patterns of protein-coding and lncRNA gene expression in a collection of 268 RNA-seq samples, derived from tumor and adjacent tissue biopsies from 114 HCC patients (Figure 1a, Supplementary Table 1). This cohort comprises patients with different underlying diseases, including hepatitis B and hepatitis C, alcoholic or non-alcoholic liver diseases and cirrhosis (Supplementary Table 1). The Edmondson-Steiner differentiation grade was recorded for each tumor sample (Supplementary Table 1). Biopsies were performed during the diagnostic work-up of patients before therapy, and in 3 patients with HCC recurrence after tumor resection (Supplementary Table 1). Our transcriptome data is not restricted to poly-adenylated RNAs (Methods) and may thus better reflect the behavior of lncRNA transcripts, which are inefficiently or not at all poly-adenylated (Schlackow *et al*, 2017). With this dataset, we could analyze the expression patterns of 19,465 protein-coding genes and 18,866 lncRNAs, including 7,959 lncRNAs detected *de novo* using our RNA-seq data (Methods, Supplementary Dataset 1).

**Figure 1.**
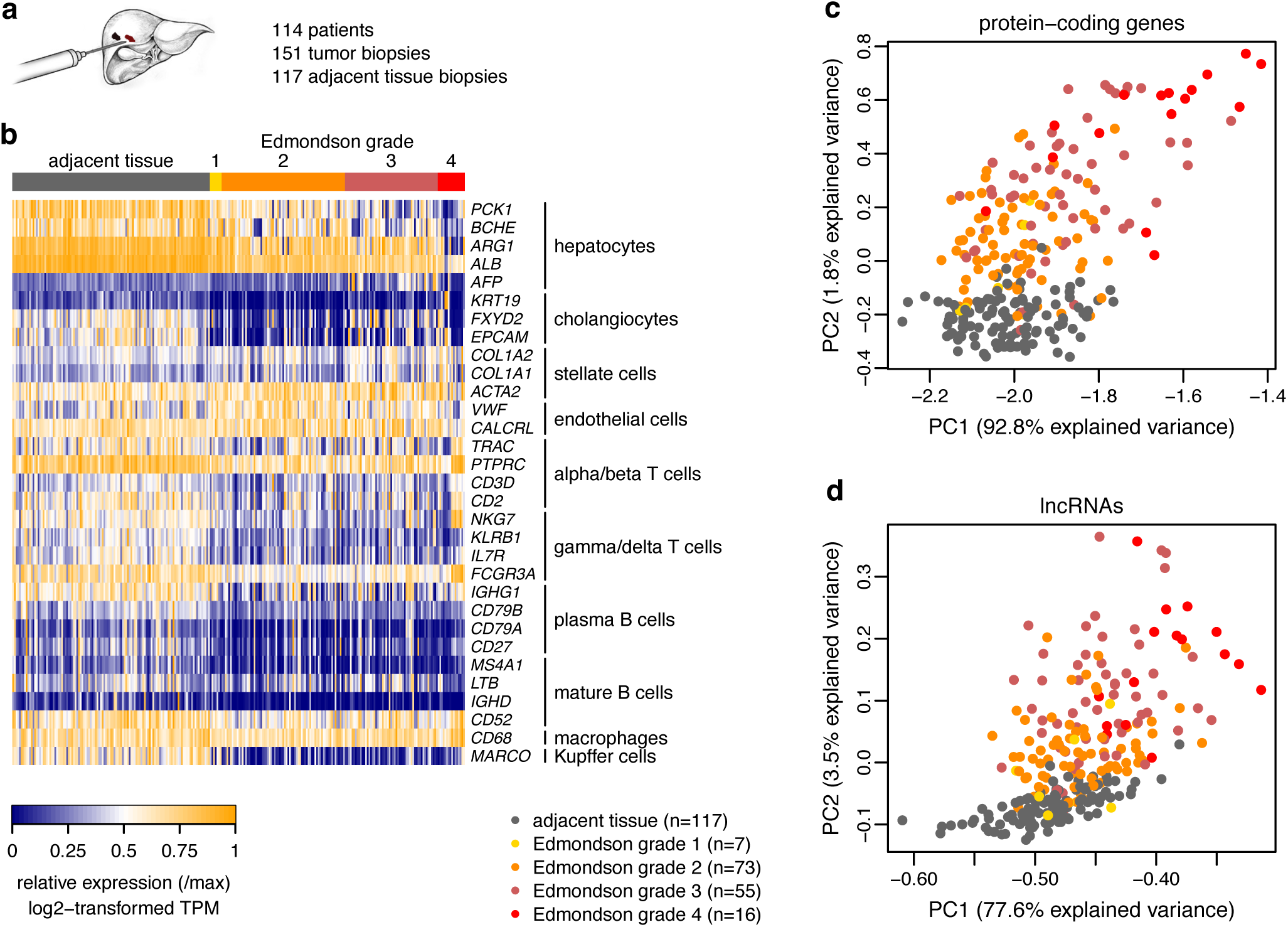
Global expression patterns in HCC tumors and adjacent tissue samples. a. Numbers of patients and RNA-seq samples included in our study. All samples are derived from pre-treatment biopsies. b. Heatmap representing relative expression levels (log2-transformed TPM values, divided by the maximum value across samples), for 36 markers of the most common cell types in the healthy liver (MacParland *et al*, 2018). c. Scatter plot representing the first factorial map of a principal component analysis, performed on log2-transformed TPM values for protein-coding genes. Each dot represents one sample. Colors represent sample types (adjacent tissue in grey, tumor samples colored according to the Edmonson-Steiner grade). d. Same as c, for lncRNAs.

### Global trends of gene expression variation in HCC tumors and adjacent tissue samples

We first aimed to evaluate broad patterns of gene expression variation among tumor and adjacent tissue samples. To get a glimpse of the cellular composition changes that take place in cancer tissue, we analyzed the expression patterns of liver cell type markers (Supplementary Table 2), obtained from single cell transcriptomics data (MacParland *et al*, 2018). As expected, many of these markers display striking differences between tumor and adjacent tissue samples, as well as among degrees of tumor differentiation (Figure 1b). Hepatocyte markers (*PCK1, BCHE, ARG1, ALB*) are low in samples derived from Edmondson-Steiner grade 4 tumors (Figure 1b). Immune cell markers (e.g., T cell markers *PTPRC, NKG7, FCGR3A* or macrophage markers *CD52* and *CD68*) are generally expressed at lower levels in tumor samples than in the adjacent tissue (Figure 1b). Overall, these patterns confirm that the cellular environment is substantially different in HCC tumors compared to the adjacent tissue, but also that there is considerable heterogeneity among tumors.

The molecular heterogeneity of HCC tumors is well illustrated by principal component analyses (PCA) performed on protein-coding and lncRNA genes (Methods, Figure 1c,d). However, although there is substantial variation among tumor samples, this gene expression map is consistent with the histological classification. For both categories of genes the first axis of the PCA separates samples with the highest Edmondson-Steiner grades and samples from less advanced tumors and adjacent tissues (Figure 1c,d, Supplementary Figure 1a-d). The second axis forms a gradient from adjacent tissue to the highest Edmondson-Steiner grades (Figure 1c,d, Supplementary Figure 1a-d). Notably, paired biopsies do not cluster on the first factorial map of the gene expression PCA, despite their shared genetic background. We validated the sample pairing by evaluating the presence of shared alleles in exonic single nucleotide polymorphisms that were reliably detected with our RNA-seq data (Methods). As expected, samples stemming from the same patient are genetically very similar, in contrast to samples derived from different patients (Supplementary Figure 1e).

In HCC, lncRNAs follow previously reported patterns: they are generally weakly expressed and are thus detected in fewer samples than protein-coding genes (Supplementary Figure 2). This trend is even stronger for *de novo* annotated lncRNAs (Supplementary Figure 2).

### Differential expression of protein-coding genes and lncRNAs in HCC

We next tested for differential expression (DE) between paired tumor and adjacent tissue biopsies and among tumors with different Edmondson-Steiner grades (Supplementary Table 3, Methods). We selected differentially expressed genes with a minimum fold change of 1.5 and maximum false discovery rate (FDR) of 1%. With these stringent settings, we found that 4,100 (21%) protein-coding genes and 3,315 (18%) lncRNAs were differentially expressed between tumor and adjacent tissue biopsies. When comparing tumor samples grouped by Edmondson-Steiner grade (grades 1 and 2 *vs.* grades 3 and 4), 2,537 (13%) protein-coding genes and 2,065 (11%) lncRNAs were significantly differentially expressed. The distribution of expression fold changes differs between the two categories of genes, with stronger positive fold changes for lncRNAs for the latter analysis (Figure 2). Genes that were up-regulated in tumors compared to adjacent tissues or in tumor samples with higher Edmondson-Steiner grades were enriched in processes related to the cell cycle, to chromosome organization but also to embryonic development (Figure 2, Supplementary Table 4). In contrast, downregulated genes were enriched in metabolic processes characteristic of the healthy liver (Figure 2, Supplementary Table 4). In addition, genes involved in immune response and in cell adhesion are down-regulated in tumor samples compared to the adjacent tissue (Figure 2a, Supplementary Table 4). There is substantial overlap between the sets of genes that are differentially expressed in the two comparisons, with consistent directions of change, for both protein-coding genes and lncRNAs (Supplementary Figure 3a,b).

**Figure 2.**
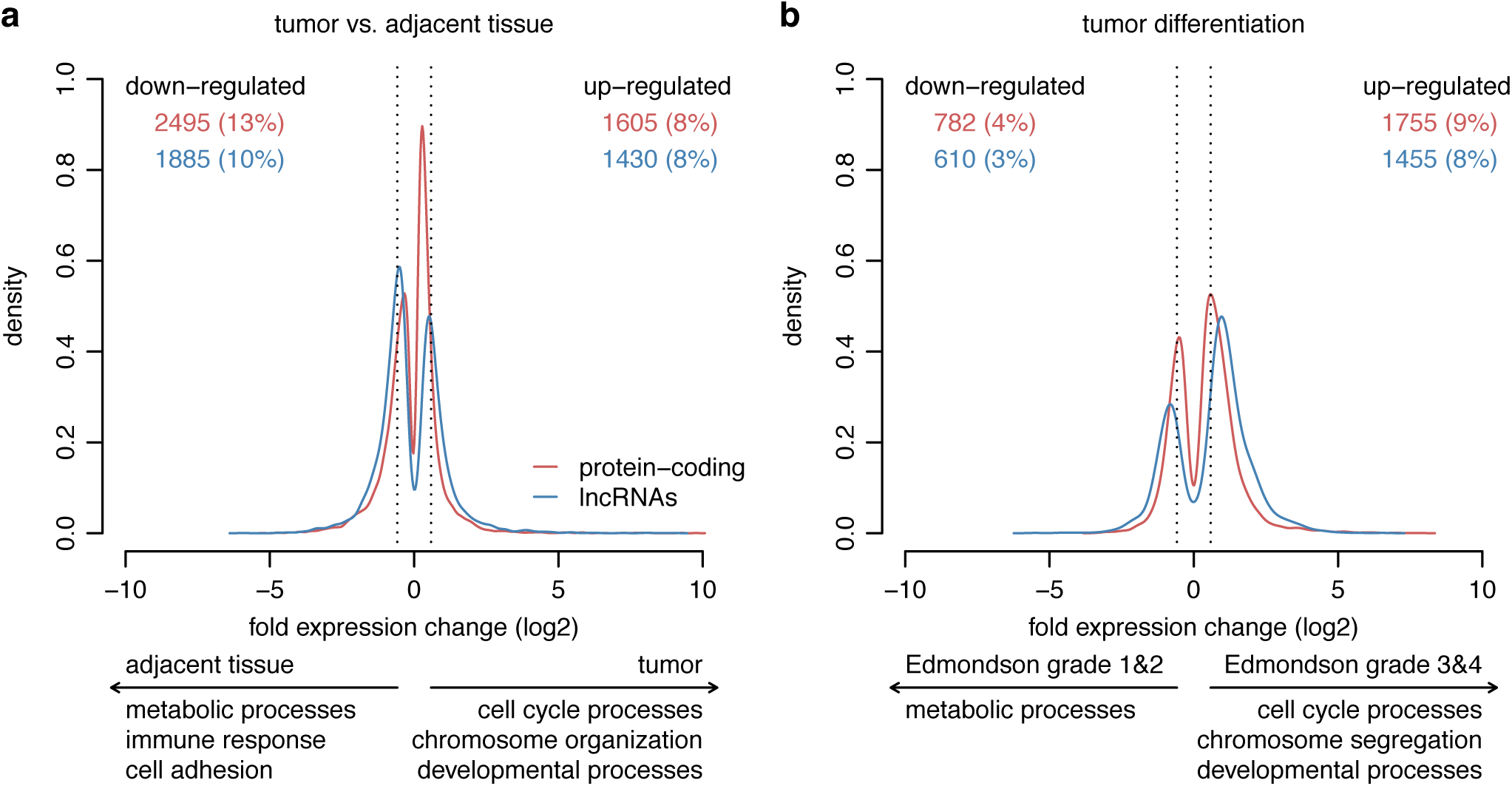
Differentially expressed genes between HCC tumors and adjacent tissue samples. a. Density plot of the log2 fold expression change, for genes that are significantly differentially expressed (maximum FDR 0.01) between paired tumor and adjacent tissue samples (Methods). Red: protein-coding genes; blue: lncRNAs. The dotted vertical lines mark an expression change threshold of 1.5. The numbers of genes that pass the FDR and minimum fold change thresholds are shown at the top of the plot. The main enriched gene ontology categories for up-regulated and down-regulated genes are shown below the plot (Methods). b. Same as a, for the analysis comparing tumor samples with different stages (Edmondson-Steiner grades 1 and 2 vs. Edmondson-Steiner groups 3 and 4).

As expected given their involvement in essential cell cycle processes, protein-coding genes that are up-regulated in tumors compared to the adjacent tissue or in the tumors with the highest Edmondson-Steiner grades had significantly higher levels of evolutionary sequence conservation than down-regulated genes (Wilcoxon rank sum test p-value < 1e-10 for the first DE analysis, p-value 0.006 for the second DE analysis, Supplementary Figure 3c). For lncRNAs, the increase in sequence conservation is only observed for those that are up-regulated in the tumors compared to the adjacent tissue (Wilcoxon rank sum test, p-value 3.7 e-4, Supplementary Figure 3d). In contrast, lncRNAs that are up-regulated in tumors with higher Edmondson-Steiner grades have slightly lower conservation scores than down-regulated lncRNAs (Wilcoxon rank sum test, p-value 0.03, Supplementary Figure 3d).

We next verified whether the DE protein-coding genes and lncRNAs were isolated or clustered in the genome. To do this, for each DE gene (defined as above) we verified the DE status for neighboring genes, within a 50 kilobases (kb) window (Methods). We find that the proportion of DE genes that have a DE neighbor with the same expression change direction is significantly higher than expected by chance, for both protein-coding genes and lncRNAs (randomization test, p-value < 0.01, Supplementary Figure 4, Methods). In contrast, pairs of neighboring genes with opposite DE orientation are significantly less frequent than expected by chance (randomization test, p-value < 0.01, Supplementary Figure 4). This pattern is observed for both protein-coding and lncRNA genes and for both differential expression tests.

Finally, we also assessed the effect of other factors (namely, underlying liver disease, presence of cirrhosis, sex of the patients) on gene expression patterns in HCC tumors. In contrast with the large numbers of DE genes observed for the two comparisons described above, only between 36 and 509 genes were significantly DE depending on one of these factors (maximum FDR 0.01, minimum fold expression change 1.5, Supplementary Dataset 3). For the comparison between sexes, 180 genes were significantly DE, with the strongest fold changes observed for genes located on sex chromosomes (Supplementary Dataset 3).

### Expression patterns of prominent HCC-associated lncRNAs

We next aimed to evaluate the behavior of the most prominent HCC-associated lncRNAs in our gene expression dataset. We performed a PubMed search with the key word “hepatocellular carcinoma” in the article title, and parsed the abstracts of the resulting articles to retrieve gene names or an unambiguous mention of lncRNAs as a class (Methods). We found that the proportion of all HCC publications that mention lncRNAs increased rapidly in the past decade, from 0 in 2010 to 6.3% in 2019 (Supplementary Figure 5a). In total, we could find unambiguous citations for 262 lncRNAs, 160 (61%) of which were only mentioned in one article (Supplementary Table 5, Supplementary Figure 5b). Only 29 lncRNAs were associated with HCC in 5 or more articles. Expectedly, at the top of the list of highly-cited lncRNAs can be found transcripts that are well known from other biological contexts, such as *MALAT1* (Ji *et al*, 2003, 1), *H19* (Bartolomei *et al*, 1991), *HOTAIR* (Rinn *et al*, 2007) and *NEAT1* (Hutchinson *et al*, 2007). The 5^th^ highest-cited lncRNA is *HULC*, which was initially described in the HCC context (Panzitt *et al*, 2007). Among the 262 HCC-associated lncRNAs, 98 (37%) were significantly DE (maximum FDR 0.01 and minimum fold change 1.5) between tumor and adjacent tissues, and 57 (22%) were significantly DE between tumor samples with Edmondson-Steiner grades 1 and 2 and tumor samples with Edmondson-Steiner grades 3 and 4. These proportions are significantly higher than those observed for lncRNAs that are not cited in the literature (17% and 11%, respectively, Chi-square test, p-value < 1e-10). In total, 128 (49%) of the HCC-associated lncRNAs were significantly DE in at least one of the tests; this proportion reached 81% with low stringency criteria (maximum FDR 0.1, no minimum fold change).

We next examined the expression patterns of the 29 lncRNAs that were cited at least 5 times in association with HCC (Figure 3). For this analysis, we set the maximum FDR at 0.01 as described above, but we did not require a minimum fold expression change, to increase our sensitivity. The great majority (90%) of these lncRNAs were significantly DE between tumors and adjacent tissues, and 14 (48%) of them were also significantly DE between highly differentiated (Edmonson-Steiner grades 1 and 2) and poorly differentiated tumors (grades 3 and 4). However, we observed several unexpected patterns among the best studied lncRNAs. First, *MALAT1* was not significantly DE in neither one of the two analyses (Figure 3), despite previous reports indicating its up-regulation in HCC tumors compared to adjacent tissues (Lin *et al*, 2007; Lai *et al*, 2012). Importantly, this is not due to a lack of statistical power or due to noisy expression, as *MALAT1* was expressed at high levels in all samples (Figure 3d). Second, *HOTAIR* was overall very weakly expressed and not significantly DE in neither of the two tests (FDR 0.046, Edmondson grades 1&2 against 3&4). Third, *NEAT1* was weakly but significantly down-regulated in tumors compared to adjacent tissues, despite previous evidence for up-regulation (Kou *et al*, 2020). For *HULC* (Panzitt *et al*, 2007), we confirmed the previously reported up-regulation in tumor samples, but surprisingly, we found that it displayed lower expression levels in advanced-stage tumors (Figure 3). In some cases, the results could be explained by the distribution of tumor differentiation degrees among the tumor samples. For example, *UCA1* is overall down-regulated in tumors compared to the adjacent tissue, contrary to what was previously reported (Wang *et al*, 2015), but is expressed at higher levels in samples with Edmondson-Steiner grades 3 and 4 (Figure 3). Some of the inconsistencies observed between our DE analyses and previous reports, for the best-studied HCC-associated lncRNAs, may also come from the distribution of patient characteristics, for example underlying liver diseases, genetic background etc. However, out of the 29 tested lncRNAs none showed significant expression differences between patients with different underlying diseases (Supplementary Dataset 3). Only *XIST* was differently expressed between sexes (Supplementary Dataset 3). We also did not observe any significant difference between patients with or without cirrhosis (Supplementary Dataset 3).

**Figure 3.**
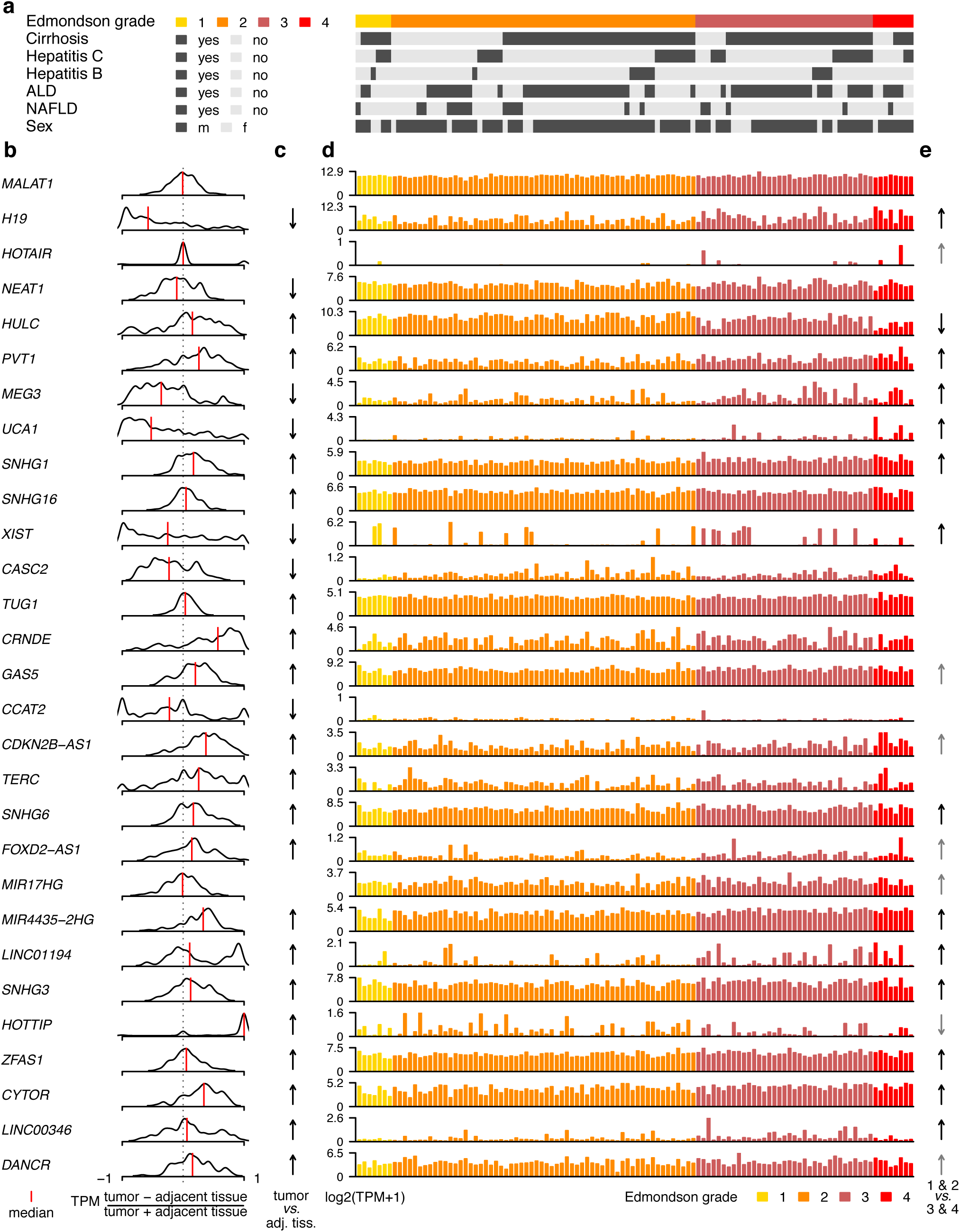
Differential expression patterns for prominent HCC-associated lncRNAs. a. Distribution of patient characteristics for the 151 tumor samples analyzed in this study. ALD: alcoholic liver disease. b. Distribution of the difference in expression levels between tumor and adjacent tissue samples, across patients, for the 29 lncRNAs that are cited in at least 5 HCC publications. The black line shows a density plot of the ratio (TPM tumor – TPM adjacent tissue)/(TPM tumor + TPM adjacent tissue), computed for each patient. Only samples used for the differential expression analyses were considered. The vertical red line represents the median value. c. Presence/absence and direction of significant expression changes between paired tumor and adjacent tissue biopsies. Upward arrows indicate up-regulation in tumor samples, downward arrows indicate down-regulation in tumor samples, with a maximum false FDR of 0.01 (no fold change requirement). Gray arrows represent marginally significant changes (FDR < 0.1, no fold change requirement). d. Expression levels (log2-transformed TPM) for tumor samples, for the 29 lncRNAs that are cited in at least 5 HCC publications. Samples are colored depending on the Edmondson-Steiner grade. e. Same as c, for the differential expression analysis comparing tumor samples with Edmondson-Steiner grades 3 and 4, *versus* tumor samples with Edmondson-Steiner grades 1 and 2.

### Increased repetitive sequence content in HCC-upregulated lncRNAs

We next wanted to assess the genomic features of the lncRNAs that are significantly differentially expressed in the two analyses described above. It was previously reported that transposable elements that are repressed in healthy tissues can become active in cancer cells (Burns, 2017). We thus analyzed the repetitive sequence content of differentially expressed lncRNAs (Supplementary Table 6, Supplementary Dataset 4, Methods). The fraction of exonic sequence covered by repeats was significantly higher for lncRNAs that were up-regulated in tumors compared to adjacent tissues (median value 47%) than for down-regulated lncRNAs (median value 40%, Wilcoxon rank sum test, p-value < 1e-10, Figure 4a). Likewise, in the DE analysis comparing tumor samples with different Edmondson-Steiner grades, up-regulated lncRNAs had significantly higher repetitive sequence content (median 48%) than down-regulated lncRNAs (median value 43%, Wilcoxon rank sum test, p-value 1e-6, Figure 4a). For protein-coding genes, the opposite trend was observed, with higher repetitive sequence contents for down-regulated genes, in both DE analyses (Figure 4a). Among the most abundant classes of repetitive elements, we found that this pattern was the strongest for satellite repeats: for both DE analyses, up-regulated lncRNAs overlap significantly more frequently with satellite repeats than down-regulated lncRNAs (Chi-square test, p-value 1e-4 for the first DE analysis, p-value 8e-5 for the second DE analysis, Figure 4b). Confirming this observation, we found that lncRNAs that overlapped with satellite repeats had significantly higher fold expression changes than lncRNAs without satellite repeats, for both DE analyses (Wilcoxon rank sum test p-value 2e-6 for the first DE analysis, 0.02 for the second DE analysis, Figure 4c). We also observed significantly higher fractions of exonic overlap with LTR repeats for lncRNAs that are up-regulated in tumors with high Edmondson-Steiner grades, compared to down-regulated lncRNAs (Supplementary Figure 6). However, for this repeat class there was no significant difference between lncRNAs that are up-regulated or down-regulated between tumors and adjacent tissues (Supplementary Figure 6).

**Figure 4.**
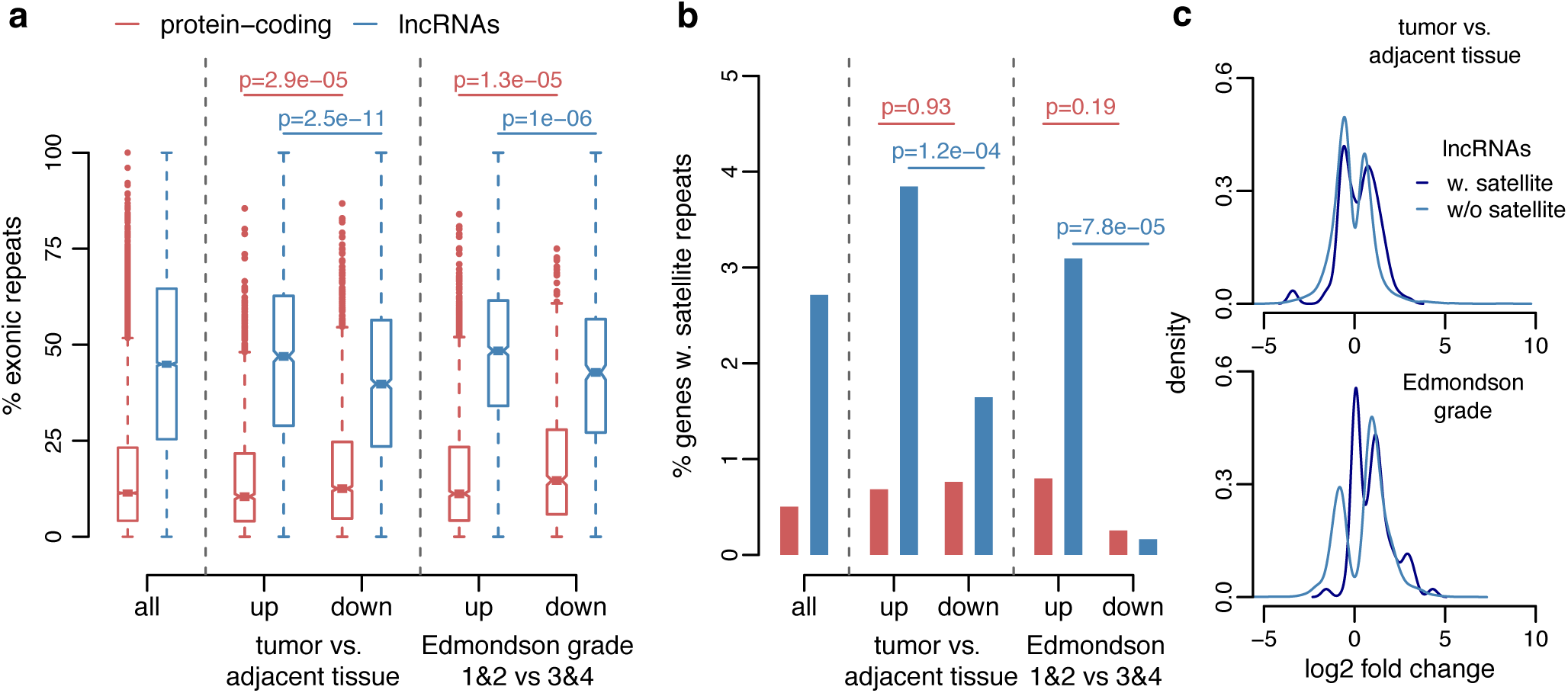
Over-representation of satellite repeats among tumor-upregulated lncRNAs. a. Boxplots of the percentage of exonic sequences covered by repetitive sequences, for protein-coding genes (red) and lncRNAs (blue). We display separately genes that are differentially expressed (maximum FDR 0.01, minimum fold expression change 1.5) in tumors compared to adjacent tissues, and in tumors with Edmondson-Steiner grades 3 and 4 compared to tumors with Edmondson-Steiner grades 1 and 2. Horizontal segments represent median values; notches represent 95% confidence intervals for the median; dashed segments extend to 1.5 times the inter-quartile range. b. Percentage of genes that have exonic overlap with satellite repeats, for protein coding genes (red) and lncRNAs (blue). As in a, we display separately genes that show significant expression differences in our two main DE analyses. c. Distribution of the log2 fold expression changes in our two main DE analyses, for lncRNAs that overlap with satellite repeats (dark blue) or not (light blue). Only lncRNAs that are show significant differences (maximum FDR 0.01, no minimum fold change requirement) are shown.

### Up-regulation of centromeric non-coding RNAs and centromeric proteins in HCC

Satellite repeats are a major functional component of centromeres (Hartley & O’Neill, 2019). Following our observation that lncRNAs that overlap with satellite repeats tend to be expressed at higher levels in tumors than in normal tissues, and in particular in advanced-stage tumors, we performed a more direct examination of transcription in centromeric regions. As these highly repetitive sequences can be difficult to capture with next generation sequencing approaches, we first determined the centromeric regions that are mappable with our RNA-seq data – that is, to which sequencing reads can be attributed unambiguously (Methods). With the exception of the Y chromosome, which had a mappable length of 222 kb, all centromeric regions had mappable lengths comprised between 1.2 Mb and 5.2 Mb (Supplementary Figure 7a). We found 752 transcribed loci in centromeric regions, all but one detected *de novo* with our RNA-seq data (Supplementary Dataset 5). In general, we could detect at most 10 centromeric transcribed loci *per* chromosome (Supplementary Figure 7b). However, we found large numbers of transcripts on chromosomes 2 and 18 (173 and 395 transcribed loci, respectively), as well as on chromosomes 1 and 19 (31 and 75 transcribed loci, respectively). With the exception of an Ensembl-annotated pseudogene, these transcripts were classified as non-coding, but only 243 passed all lncRNA filtering criteria (Supplementary Dataset 5). The other non-coding transcripts were generally rejected from the lncRNA dataset because they were too short (38% of the cases), they overlapped with unmappable regions (14%), they had insufficient read coverage (4%), or because of a combination of these criteria.

We evaluated the abundance of centromeric transcripts by counting unambiguously mapped RNA-seq for each chromosome and strand, normalized by dividing by the total unique read count attributed to genes, for each sample (Methods, Supplementary Dataset 5). Most centromeric RNA-seq reads were derived from chromosome 2, followed by chromosome 1 and 19 (Figure 5a). Chromosome 2 also stood out with respect to the differences between tumor and adjacent tissue samples: on the reverse DNA strand, 94 patients (85%) had higher transcript levels in tumors than in adjacent tissue samples (Figure 5b). We note that transcription is not restricted to well-defined loci, but covers the entire centromeric region (Figure 5c).

**Figure 5.**
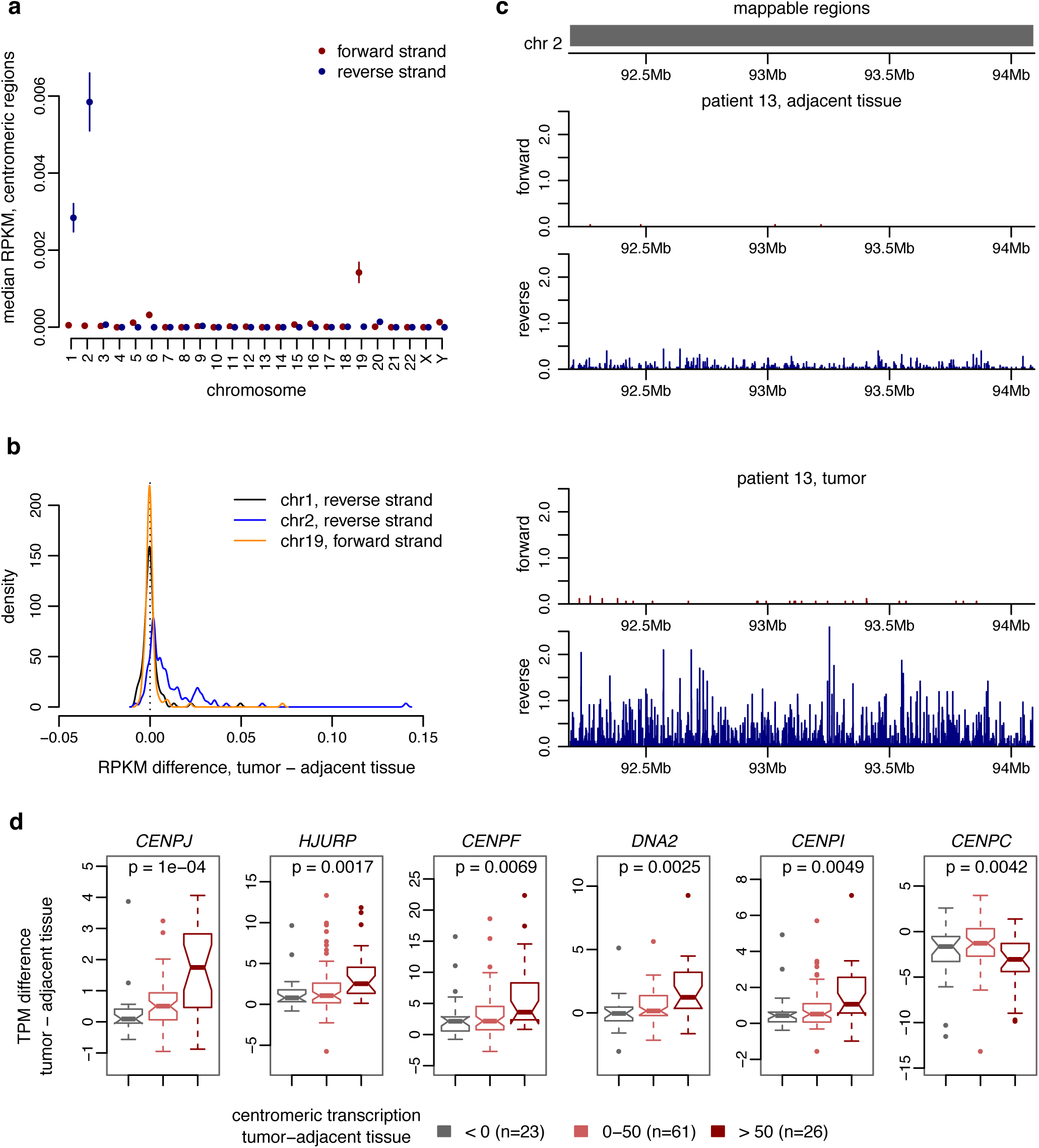
Increased centromeric transcription in tumors compared to adjacent tissue samples. a. Dot chart representing the median normalized expression levels (RPKM) for centromeric regions, across samples, for each chromosome and strand. Red: transcripts on the forward DNA strand, blue: reverse strand. The bars represent the 95% confidence intervals. b. Density plot of the RPKM difference between tumor and adjacent tissue, across patients, for the three chromosome/strand combinations with highest RPKM levels (chromosome 2 reverse, 1 reverse and 19 forward strand). c. Top: representation of the regions considered to be unambiguously mappable (Methods), for the chromosome 2 centromere. Next panels: unique read coverage distribution on the chromosome 2 centromere, forward and reverse strands, for one patient (identifier 13). The read coverage was normalized for each sample based on the number of million mapped reads attributed to genes. d. Boxplots representing the distribution of gene TPM differences between tumor and adjacent tissues, for three classes of patients defined based on the degree of centromeric transcript “activation” in tumors compared to adjacent tissues. The first class comprises 23 patients for which the difference in RPKM values for total centromeric transcripts between tumors and adjacent tissues is below 0 RPKM; the second class comprises 61 patients for which the difference is between 0 and 50 RPKM; the third class comprises 26 patients for which the difference is above 50 RPKM. We display the 6 genes mentioned in the text: *CENPJ, HJURP, CENPF, DNA2, CENPI, CENPC.* P-values correspond to Kruskal-Wallis non-parametric tests, for differences among the three classes of patients.

The degree of centromere transcript activation in tumor samples compared to adjacent tissue samples varies considerably among patients (Figure 5b). To evaluate the determinants of centromeric transcription variation, we analyzed the association between protein-coding gene differential expression and centromeric transcript differential expression, across patients. Specifically, for each patient, we computed the difference in TPM levels between tumors and adjacent tissues, for each protein-coding gene; we also computed the difference in total centromeric RPKM levels between tumors and adjacent tissues, and we correlated the two sets of values across patients. Genes involved in mitotic cell cycle processes were often positively associated with centromeric transcript activation levels (Supplementary Table 7, gene ontology enrichment analysis presented in Supplementary Dataset 5). Among the genes with the highest positive correlations with centromeric transcript activation levels were several genes encoding centromeric proteins (*CENPJ*, *CENPF* and *CENPI*), the CENPA chaperone *HJURP* (Hori *et al*, 2020), the *DNA2* nuclease/helicase that promotes centromeric DNA replication (Li *et al*, 2018), etc. (Figure 5d, Supplementary Table 7). Interestingly, *CENPC*, which is thought to repress alpha-satellite RNA levels (Bury *et al*, 2020), was negatively associated with centromeric transcript activation levels (Figure 5d, Supplementary Table 7).

In addition to increased levels of centromeric non-coding RNAs in tumors, we also observed a strong tendency for up-regulation for centromeric proteins (Supplementary Table 8). Out of 25 protein-coding genes annotated in Ensembl as “centromere proteins”, 20 (80%) were up-regulated in the tumors compared to adjacent tissue and 13 (52%) were up-regulated in tumors with Edmondson-Steiner grades 3&4 compared to tumors with Edmondson-Steiner grades 1&2 (maximum FDR 0.01). At the top of the list, the genes coding for the histone variant *CENPA* and for centromeric protein F (*CENPF*) were more than 4-fold over-expressed in tumors compared to adjacent tissues (Supplementary Table 8). Confirming our previous analysis, we also observed that *CENPC* was down-regulated in tumors compared with adjacent tissues and in advanced-stage tumors compared to early-stage tumors (Supplementary Table 8).

## Discussion

### Protein-coding gene and lncRNA expression patterns in HCC

With this analysis, our first aim was to investigate the gene expression alterations that characterize HCC tumors. Compared to the numerous transcriptome collections that were previously published in the HCC field, our dataset has the advantage of including a large number of paired tumor and adjacent tissue samples, comprising a total of 268 samples from 114 patients. Importantly, the samples analyzed here are derived from biopsies, which are likely to better reflect the situation *in vivo,* because they are devoid of changes induced by hypoxia and hypoglycemia that occur in surgical resection specimens as a consequence of segmental blood vessel occlusions during the operation. Moreover, our data includes both poly-adenylated and non-poly-adenylated RNA species, which makes it better suited for the study of inefficiently poly-adenylated lncRNAs (Schlackow *et al*, 2017).

We first explored the broad patterns of gene expression variation in our tumor and adjacent tissue samples. By analyzing the expression patterns of molecular markers for the most common cell types in the healthy liver (MacParland *et al*, 2018), we confirmed that HCC tumors have very different cellular environments compared to adjacent tissue samples (Figure 1). In particular, immune cell populations appear to be diminished in the majority of tumors, (Figure 1). Although these patterns are evidently better investigated with single-cell RNA-seq data, these results confirm that our transcriptome collection reflects the cellular composition changes that define the “tumor microenvironment” (Hanahan & Weinberg, 2011).

As expected, we found that gene expression patterns are in good agreement with the histological classification of the tumor samples. For both protein-coding genes and lncRNAs, tumor samples with Edmondson-Steiner grades 3 and 4 stand out from tumors with lower grades and from adjacent tissue samples (Figure 1, Supplementary Figures 1). Other factors, such as the underlying liver disease, the sex or the age of the patients, have comparatively little effect on the overall gene expression variation. We thus focused on the protein-coding genes and lncRNAs that are differentially expressed between paired tumor and adjacent tissue samples, or between poorly differentiated tumors (Edmondson-Steiner grades 3 and 4) and highly differentiated tumors (Edmondson-Steiner grades 1 and 2). We observed an over-representation of biological processes associated with the cell cycle among genes that are up-regulated in the tumors (Figure 2), which is expected given that cancer cells are rapidly proliferating. Conversely, genes involved in the metabolic processes performed by the healthy liver or in immune response tend to be down-regulated in the tumors (Figure 2).

Both protein-coding genes and lncRNAs contribute to the differential gene expression patterns observed in HCC tumors (Figure 2). Differentially expressed protein-coding genes and lncRNAs share many characteristics. For example, for both gene categories, we found that genes that are up-regulated in tumors compared to adjacent tissue samples have significantly higher levels of evolutionary sequence conservation than genes with the opposite expression change (Supplementary Figure 3). This observation is consistent with the enrichment of cell cycle functions among protein-coding genes that are up-regulated in the tumors, as these genes have essential biological roles and are thus under strong constraint during evolution. The increase in sequence conservation for lncRNAs that are up-regulated in tumors suggests that these lncRNAs may also participate in essential cellular functions and contribute to cellular proliferation. Another shared feature between protein-coding genes and lncRNA is the presence of spatial clustering: differentially expressed genes are found in close proximity to other differentially expressed genes with the same expression change direction significantly more often than expected by chance (Supplementary Figure 4). This observation may be explained by a tendency for co-regulation of neighboring lncRNA and protein-coding genes, or may reflect the presence of large-scale structural variations (rearrangements, duplication and deletions) in cancer cells, which can affect the expression patterns of multiple neighboring genes (Spielmann *et al*, 2018). This finding also underlines the importance of evaluating the broader genomic context when aiming to select candidate oncogenes, tumor suppressors, or biomarkers: the most biologically relevant gene may be the neighbor of the gene initially selected for validation.

### Limited reproducibility of differential expression patterns for HCC-associated lncRNAs

In the past decade, the number of publications that discuss lncRNAs in the context of hepatocellular carcinoma has increased exponentially (Supplementary Figure 5). LncRNAs are often proposed as promising oncogenes or tumor suppressors, based on their patterns of expression in tumors and healthy tissues. However, lncRNAs are weakly expressed and are generally highly variable among tissues, cell types or individuals (Kornienko *et al*, 2016). Thus, it is not clear to what extent the lncRNA expression patterns previously reported in the HCC literature are reproducible with independent datasets. Here, we evaluated the expression patterns of lncRNAs that were previously associated with HCC in our transcriptome collection. The majority of these lncRNAs were strongly differentially expressed between paired tumors and adjacent tissue samples or between groups of tumors with high or low differentiation. However, we are still far from confirming differential expression patterns for all HCC-associated lncRNAs, even when lowering the stringency of our criteria. Even when evaluating the most prominent HCC-associated lncRNAs, which were cited by at least 5 publications, we could not always recover the previously reported differential expression observations. This was the case even for the three lncRNAs that were most frequently associated with HCC in the literature: MALAT1, H19 and HOTAIR. The biological roles of these lncRNAs in cancer were already controversial. For example, a recent study showed that MALAT1 suppresses metastasis in breast cancer (Kim *et al*, 2018, 1), contrary to previous reports which proposed that this lncRNA promotes metastasis (Arun *et al*, 2016). Likewise, H19 was alternatively proposed as an oncogene (Matouk *et al*, 2007) or as a tumor suppressor (Yoshimizu *et al*, 2008). For HOTAIR, its role as a metastasis-promoting factor appears to be accepted in the literature (Gupta *et al*, 2010). However, we note that the initially proposed function for this lncRNA, namely a role in the regulation of *HOXD* genes during embryonic development (Rinn *et al*, 2007), was refuted *in vivo* (Amândio *et al*, 2016). These examples illustrate the frailty of some of the claims that are recurrently put forward regarding lncRNA functions, in cancer or in other biological contexts, and again highlight the caution that should be exercised when investigating lncRNAs.

### Activated transcription of centromeric satellite repeats in HCC tumors

Transcriptome comparisons in HCC cohorts or in other cancer types generally aim to select candidate oncogenes, tumor suppressors or biomarkers, that should be further verified experimentally. As extensive functional validations were outside of the scope of our study, we chose instead to analyze the genomic characteristics of the lncRNAs that were differentially expressed in HCC tumors. We were thus able to detect an increase in the repetitive sequence content of lncRNAs that were up-regulated in tumors compared to adjacent tissues, as well as in poorly differentiated tumors compared to early stage tumors (Figure 4). Repetitive sequences make up roughly half of the human genome (Lander *et al*, 2001). The high repeat fraction observed for lncRNA exons, which is more than triple the fraction observed for protein-coding gene exons (Figure 4), is likely due to the weak selective pressures that act on these loci (Darbellay & Necsulea, 2020). However, the increase in repetitive sequence content for tumor-upregulated lncRNAs cannot simply be explained by a lower proportion of functionally constrained loci; on the contrary, average sequence conservation scores are higher for tumor-upregulated lncRNAs than for tumor-downregulated lncRNAs (Supplementary Figure 3). Moreover, we found that the over-representation of repetitive sequences in upregulated lncRNAs does not affect all classes of repeats, but is strongest for satellite repeats (Figure 5). This class of repeats is a major functional component of centromeres (Hartley & O’Neill, 2019).

Although centromeres were initially thought to be transcriptionally inert, it is now known that they are transcribed into non-coding RNAs, which associate with centromeric chromatin and potentially participate in kinetochore formation (Talbert & Henikoff, 2018). However, these non-coding RNAs are generally weakly transcribed, and higher expression levels can lead to impaired centromeric function (Bouzinba-Segard *et al*, 2006). Overexpression of centromeric non-coding RNAs was previously reported in pancreatic cancers and in other types of epithelial cancers (Ting *et al*, 2011). In mouse models of pancreatic cancers, it was shown that overexpression of centromeric satellite repeats leads to increased DNA damage and chromosomal instability, thereby accelerating tumor formation (Kishikawa *et al*, 2016, 2018). Here, we reveal that centromeric non-coding RNAs are also aberrantly overexpressed in HCC. This finding is supported by several lines of evidence. First, we showed that satellite repeats, which are characteristic of centromeric regions, are over-represented in the exonic regions of tumor-upregulated lncRNAs. Second, we directly quantified centromeric transcription, by evaluating regions to which RNA-seq reads can be unambiguously attributed, despite the repetitive sequence context. We thus showed that transcription stems from the entire length of centromeric regions, rather than from well-defined non-coding RNA loci. Interestingly, all chromosomes are not equal with respect to detectable centromeric transcription. The centromere of chromosome 2 appears to be transcriptionally active in tumor samples for the majority of patients (Figure 5). The mechanisms that underlie this over-representation of chromosome 2 are unclear. This chromosome has a particular evolutionary history: it is derived from a chromosome fusion event, which occurred after the divergence of human and chimpanzee and which led to the loss of one of the two ancient centromeres (Chiatante *et al*, 2017). Although we verified that the over-representation of chromosome 2 is not simply due to a better mappability of satellite repeats (Supplementary Figure 7), we cannot exclude other technical issues that prevent us from detecting these highly repetitive transcripts from other chromosomes.

The levels of centromeric non-coding RNA transcription were previously found to vary during the cell cycle in mouse, with a peak in the G2/M phase (Ferri *et al*, 2009). Thus, our findings may be partially explained by an over-representation of cells in the G2/M phase in tumor samples compared to the adjacent tissue, expected given that cancerous cells are rapidly proliferating. Indeed, our analysis revealed that genes involved in mitotic cell cycle processes were positively associated with centromeric transcript up-regulation levels, across patients. This included several genes encoding centromeric proteins (*CENPJ*, *CENPF* and *CENPI*) (Figure 5d, Supplementary Table 7). Interestingly, the gene encoding centromere protein C (*CENPC*) was negatively associated with centromeric transcript up-regulation levels across patients, and was significantly down-regulated in tumors compared to adjacent tissues and in advanced stage tumors (Figure 5d, Supplementary Tables 7-8). It was recently reported that this protein acts to repress centromere-derived alpha-satellite RNA levels (Bury *et al*, 2020). This observation could thus explain the up-regulation of centromeric transcripts in tumor compared to adjacent tissue samples, which appears to occur in parallel with a down-regulation of *CENPC* expression.

We also note that the ability to detect centromeric non-coding RNAs likely depends on the methods used to generate RNA-seq data. Our transcriptome collection was generated from ribo-depleted RNA samples, without enrichment for poly-adenylated RNA species (Methods). Although it was reported that centromeric transcripts are poly-adenylated (Topp *et al*, 2004), their subsequent processing into smaller RNA molecules (Talbert & Henikoff, 2018) may lead to the loss of the polyA tail, thus hampering their detection in polyA-enriched RNA-seq data. Furthermore, our RNA-seq data consists of relatively long reads (126-136 bp), which likely increases our ability to unambiguously map RNA-seq reads to the genomic regions from which they stem, even in the case of repetitive sequences.

To our knowledge, aberrant transcription of centromeric non-coding RNAs had not been previously reported in HCC. Given that this phenomenon has been associated with tumor formation in other types of cancer (Kishikawa *et al*, 2018), our observations are highly relevant for the search for oncogenic factors driving hepatocellular carcinoma, and thus warrant further investigations.

## Methods

### Biological sample collection

The analyses presented in this manuscript were performed on carcinoma and adjacent liver tissue biopsies obtained from 114 patients. Human tissues were obtained from patients undergoing diagnostic liver biopsy at the University Hospital Basel. Written informed consent was obtained from all patients. The study was approved by the ethics committee of the northwestern part of Switzerland (Protocol Number EKNZ 2014-099). The samples analysed here were derived from pre-treatment biopsies, with the exception of 3 patients, for which samples were collected after tumor resection (Supplementary Table 1). We recorded the sex, age at the time of biopsy and underlying liver diseases for each patient (Supplementary Table 1). We also recorded the percentage of tumor tissue in the biopsies and the Edmondson-Steiner grades of the tumors (Supplementary Table 1). Multiple tumor and adjacent tissue biopsies were collected for 26 and 3 patients, respectively. In total, we analysed 268 samples, corresponding to 117 adjacent tissue and 151 tumor biopsies.

### RNA extraction and library preparation

We extracted RNA and DNA from tissue biopsies using the ZR-Duet DNA/RNA MiniPrep Plus kit (Zymo Research, catalog number D7003). We performed the in-column DNase I treatment as specified in the kit to remove residual DNA from the RNA fraction. We prepared RNA-seq libraries using the Illumina TruSeq stranded RNA protocol, without polyA selection. We depleted ribosomal RNA using the Ribo-Zero Gold kit from Illumina. We generated single-end reads, 126 or 136 nucleotides (nt) long (Supplementary Table 1).

### RNA-seq data processing

We aligned the RNA-seq reads on the genome using HISAT2 (Kim *et al*, 2015, 2) version 2.0.5. We used the primary assembly of the human genome version GRCh38 (hg38), downloaded from Ensembl (Cunningham *et al*, 2019). We built the HISAT2 genome index using additional splice site information from Ensembl release 97, as well as from the CHESS (Pertea *et al*, 2018) and MiTranscriptome (Iyer *et al*, 2015) transcript assemblies. We extracted unambiguously mapped reads based on the NH tag from HISAT2 reported alignments. To evaluate the prevalence of strand errors during library preparation, we identified introns with GT-AG and GC-AG splice sites, supported by spliced RNA-seq reads aligned on at least 8 nucleotides on each neighboring exon and with a maximum mismatch frequency of 2%. We then compared the strand inferred based on splice site information with the strand inferred based on the read alignment orientation and on the library type. All libraries had strand error rates below 2.5% (Supplementary Table 1). The presence of contradictory strand assignments was used as a red flag in our lncRNA filtering procedure (see below).

### Single nucleotide polymorphism analysis

We verified that samples derived from the same patient were correctly paired by assessing their genetic similarity, using RNA-seq information alone. To do this, we first scanned the RNA-seq alignments to detect putative single nucleotide polymorphisms (SNPs). We used a a pipeline combining tools from GATK (Van der Auwera *et al*, 2013) version 4.1.9.8 and Picard (http://broadinstitute.github.io/picard/) version 2.18.7. Briefly, we analyzed non-duplicated aligned RNA-seq reads, re-calibrated the alignment quality around known variants from dbSNP (Sherry *et al*, 2001) release 151 and called variants with a minimum base quality score threshold of 20. We combined the detected SNPs across all samples and filtered them to keep only positions found in dbSNP and in exonic regions, excluding repetitive sequences. For all resulting SNPs, we counted the number of reads supporting each allele using the ASEReadCounter tool. We kept biallelic SNPs supported by at least 50 reads. To allow for sequencing or mapping errors, SNPs were considered to be heterozygous if the estimated allele frequency was between 0.1 and 0.9, and homozygous if the allele frequency was equal to 0 or 1. For all pairs of samples, we computed the proportion of SNPs with shared alleles out of all biallelic SNPs. We compared this measure of genetic similarity between pairs of samples derived from the same patient or from different patients (Supplementary Figure 1). We also evaluated the proportion of heterozygous SNPs out of all detected SNPs on autosomes and on sex chromosomes, for each sample. We excluded the pseudo-autosomal regions from sex chromosomes. For one male patient (identifier 42), we observed high levels of heterozygosity on the X chromosome and high *Xist* expression levels, for both tumor and adjacent tissue biopsies. This patient was excluded from differential expression analyses (Supplementary Table 1).

### Evaluation of genomic DNA contamination

To assess the amount of genomic DNA contamination, we evaluated the RNA-seq read coverage on repeat-masked intergenic regions, on both forward and reverse strands. As genuinely transcribed regions are generally strongly biased in favor of one strand, we computed the number of regions that had relatively symmetric strand distribution, *i.e.* for which the absolute value of the (forward-reverse)/(forward+reverse) coverage ratio was below 0.5. We then computed for each sample the proportion of intergenic regions with symmetric coverage, out of all intergenic regions with RNA-seq coverage. We considered that samples with more than 5% symmetrically transcribed intergenic regions had significant DNA contamination. These samples were excluded from differential expression analyses (Supplementary Table 1).

### Identification of “mappable” and “unmappable” genomic regions

To determine whether RNA-seq reads can be correctly traced back to their genomic region of origin, we performed a “mappability” analysis. To do this, we generated single-end sequencing reads with the same lengths as in our data (126 and 136 nt) from sliding genomic windows with 5 nt step. Reads were generated with perfect sequence quality and no mismatches. We aligned these reads on the genome using HISAT2 with the same parameters as for the real RNA-seq data. Genomic regions to which simulated reads were mapped back unambiguously and on their entire length were said to be mappable. We defined unmappable regions by subtracting mappable intervals from full-length chromosomes.

### Transcript assembly

We performed a genome- and transcriptome-guided transcript assembly with StringTie (Pertea *et al*, 2015) release 2.1.2. We used as an input the unambiguously mapped reads obtained with HISAT2, combined across all samples. We used annotations from Ensembl (Cunningham *et al*, 2019) release 99, excluding read-through transcripts, as a guide for the assembly. We ran StringTie separately for each chromosome and strand; unassembled contigs and the mitochondrion were excluded. We filtered the StringTie output to discard artefactual antisense transcripts stemming from library preparation errors. To do this, we computed the sense and antisense exonic read coverage for each transcript and kept only those transcripts which had a sense/antisense ratio of at least 5% in at least one sample. We also removed transcripts that contained splice junctions with contradictory strand assignments based on the splice site (GT-AG or GC-AG) and on the read alignment and library type. We combined Ensembl 99 and filtered StringTie transcript annotations by adding to the Ensembl reference those *de novo* annotated transcripts which had exonic overlap with at most 1 Ensembl-annotated gene. Ensembl-annotated transcripts were not altered, with the exception of read-through transcripts (defined as transcripts that overlap with more than one multi-exonic gene), which were discarded. LncRNAs that overlapped with annotated microRNAs were annotated separately from the miRNA products.

### Protein-coding potential of newly assembled transcripts

We used the PhyloCSF (Mudge *et al*, 2019) codon substitution frequency score to evaluate the protein-coding potential of newly assembled transcripts. To do this, we overlapped exonic coordinates with protein-coding regions predicted by PhyloCSF, in all possible reading frames. Transcripts were said to be potentially protein-coding if they overlapped with a PhyloCSF protein-coding region on at least 150 nt. Due to the nature of the genetic code, some substitutions are synonymous on both DNA strands, which can generate artefactually high PhyloCSF scores on the antisense strand of protein-coding regions. We thus required that the overlap with PhyloCSF regions be higher on the sense strand than on the antisense strand of the transcripts. We also evaluated the similarity between lncRNA sequences and known proteins and protein domains, using DIAMOND (Buchfink *et al*, 2015) against SwissProt (UniProt Consortium, 2019) and Pfam (El-Gebali *et al*, 2019). We retained SwissProt entries with high confidence scores (1 to 3) and the Pfam-A subset of Pfam. We searched for hits on repeat-masked cDNA sequences with the “blastx” flavor of DIAMOND and we required a maximum e-value of 0.01. Transcripts were said to be potentially protein-coding if they had similarity with a known protein or protein domain on at least 150 nt, with at least 40% sequence identity. Genes were said to be potentially protein-coding if at least one of their isoforms was predicted as protein-coding with either method.

### LncRNA dataset

We established a lncRNA dataset by combining lncRNAs annotated in Ensembl (gene biotype “lncRNA”) and transcribed loci annotated with StringTie that passed several filters: no protein-coding potential, evaluated as described above; minimum exonic length of 200 nt for multi-exonic loci and 500 nt for mono-exonic loci; at most 5% exonic length overlap with unmappable genomic regions; no overlap with Ensembl-annotated protein-coding genes on the same strand; at least 5000 nt away from protein-coding gene exons; at most 25% exonic length overlap with RNA repeats; at most 10% exonic length overlap with retrogenes (coordinates downloaded from the UCSC Genome Browser database (Casper *et al*, 2018)). We also required transcribed loci to be supported by at least 100 RNA-seq reads. LncRNA annotations are provided in Supplementary Dataset 1 online.

### Literature search for HCC-associated lncRNAs

We searched for articles in PubMed with the key word “hepatocellular carcinoma” in the article title. We retrieved the article abstract, title, journal and publication date. We searched for gene names in the abstract, based on a list of common gene names and synonyms in the Ensembl database. We excluded gene names that were ambiguous and matched with common terms in the HCC literature (e.g., MRI, TACE, etc). We also checked if articles contained general references to lncRNAs as a class, based on the “long non-coding RNA” and “lncRNA” keywords, with spelling variations (e.g. “noncoding” instead of “non-coding”, “lincRNA” instead of “lncRNA”, etc.).

### Gene expression estimation

We evaluated gene expression values with Kallisto (Bray *et al*, 2016) release 0.46.1 (patch by P.V. to correct bootstrap estimates). We obtained effective read counts and transcript *per* million (TPM) values for each isoform and obtained gene-level TPM values using tximport (Soneson *et al*, 2015) in R. We performed an additional normalization across samples, with a previously-proposed median-scaling approach based on the 100 genes that vary least in terms of expression ranks among samples (Brawand *et al*, 2011). This approach was applied on gene-level TPM values. For most gene expression analyses, we used log2-transformed TPM values, adding an offset of 1 (TPM to log2(TPM+1)). As a control, we also evaluated unique read counts per gene using featureCounts from Rsubread (Liao *et al*, 2019). Expression estimation analyses were performed on the full set of detected transcribed loci, including protein-coding genes, lncRNAs and other types of genes. Gene expression levels are provided in Supplementary Dataset 2 online.

### Principal component analyses

We performed principal component analyses using the dudi.pca function in the ade4 library in R (Dray & Dufour, 2007). We used log2-transformed TPM levels, for all protein-coding and lncRNA genes or for each gene type separately. We enabled variable centering but not scaling and kept 5 axes.

### Differential expression analyses

We used DESeq2 release 1.28.0 and txImport (Soneson *et al*, 2015) release 1.16.1 in R to assess differential expression, based on Kallisto-estimated effective read counts *per* transcript. We performed all differential expression analyses on the combined set of protein-coding and lncRNA genes. Given that the number of biopsies varied among patients, we first selected one pair of tumor and adjacent tissue samples *per* patient, to ensure patients contributed equally to DE results. For patients where biopsies were done before and after tumor resection, we selected the biopsies obtained before resection. For one patient, an adjacent tissue biopsy was performed before onset of HCC; we excluded it from DE analyses. For all other cases where multiple biopsies were available, we selected the sample with the largest number of uniquely mapped reads for each tissue type. We tested for differential expression between pairs of tumors and adjacent tissues by fitting a model that explains gene expression variation as a function of two factors, the tissue type and the patient of origin. We then evaluated the difference between tumors and adjacent tissues with a Wald test contrasting the two tissue types and estimated the effect size with the “apeglm” shrinkage method (Zhu *et al*, 2019). We repeated this analysis separately for males and females. We also tested for differences in gene expression among tumor samples, depending on the patient sex Edmondson-Steiner grade, presence or absence of hepatitis C, hepatitis B, cirrhosis, alcoholic liver disease or non-alcoholic liver disease. To do this, we fitted an additive model including all these factors and then evaluated the effect of each factor by contrasting its levels with a Wald test, using the “apeglm” shrinkage method to estimate the effect size (Zhu *et al*, 2019). For the Edmondson-Steiner grade, we contrasted grades 1 and 2 against grades 3 and 4. We performed a preliminary test for an age effect, but as there were no significantly DE genes this factor was not included in the model. Differential expression analyses were performed only on protein-coding and lncRNA genes. Results are provided in Supplementary Dataset 3 online.

### Gene ontology analyses

We performed gene ontology enrichment analyses with GOrilla (Eden *et al*, 2009), contrasting the lists of up- or down-regulated protein-coding genes in each test with a background set consisting of protein-coding genes expressed in those samples. To define the background set, we evaluated the minimum expression level (DESeq2-normalized read counts) of differentially expressed genes and selected genes that had higher or equal expression levels. For the analysis of the association between protein-coding gene expression and centromeric transcript levels, across patients, we analyzed the gene ontology enrichment in a single ranked list, comparing genes at the top of the list (with high, positive correlation coefficients) to genes at the bottom of the list (with low, negative correlation coefficients).

### Cell type marker analyses

We analyzed the expression patterns of common markers for the most frequent cell types in the liver from a single cell RNA-seq study (MacParland *et al*, 2018). We computed a Z-score matrix from the log2-transformed normalized TPM values across samples.

### Sequence conservation analyses

We downloaded PhastCons (Siepel *et al*, 2005) sequence conservation scores, computed on a multiple genome alignment on human and 29 other mammalian species, from the UCSC Genome Browser (Casper *et al*, 2018). We computed average PhastCons scores on exonic regions and splice sites. For loci that overlapped with other genes, we also computed average scores on non-overlapping exonic regions. Results are provided in Supplementary Dataset 4 online.

### Repetitive sequence analyses

We downloaded repetitive element coordinates predicted with RepeatMasker (Smit *et al*, 2003) from the UCSC Genome Browser (Casper *et al*, 2018). We overlapped the exonic coordinates of all protein-coding and lncRNA loci with repetitive elements and we analyzed the exonic fraction covered by each repeat class. Results are provided in Supplementary Dataset 4 online.

### Centromeric transcription analyses

We downloaded centromeric region coordinates from the UCSC Genome Browser (Casper *et al*, 2018). We determined the mappable regions within each centromere as described above, by discarding regions deemed unmappable for 126 or 136 nt read lengths. We counted the number of unambiguously mapped reads that could be attributed to each mappable centromeric region, on each DNA strand, using featureCounts in the Rsubread R package (Liao *et al*, 2019). We computed normalized expression values (RPKM) by dividing the read counts by the mappable region length (expressed in kilobases) and by the number of million mapped reads, counted on the gene models annotated in Ensembl or detected with StringTie. We extracted centromeric proteins based on the “centromere” keyword in the Ensembl gene description. We used the same set of samples selected for differential expression analyses. Results are provided in Supplementary Dataset 5 online. Centromeric transcription read coverage tracks are available online for all patients: http://pbil.univ-lyon1.fr/members/necsulea/MERIC_lncRNAs/.

### Co-expression between centromeric transcript levels and protein-coding gene expression

To evaluate the determinants of centromeric transcription variation, we estimated the correlation between protein-coding gene expression and centromeric transcript levels across patients. Specifically, for each patient we estimated the difference in centromeric RPKM between tumor and adjacent tissue samples, using the samples selected for differential expression analyses. In parallel, we computed the difference in gene TPM between tumor and adjacent tissue samples, for all protein-coding genes. We then computed Spearman’s correlation coefficients for each protein-coding gene, using the values described above for all patients. Results are presented in Supplementary Table 7 and in Supplementary Dataset 5 online.

### Data and code availability

The sequencing data used in this project was submitted to the European Genome-Phenome Archive under the accession number EGAS00001004976. Supplementary datasets containing all the information needed to reproduce the results are available at the address: http://pbil.univ-lyon1.fr/members/necsulea/MERIC_lncRNAs/. Scripts are available in GitLab: https://gitlab.in2p3.fr/anamaria.necsulea/meric.

## Acknowledgements

This work was supported by European Research Council Synergy grant 609883 Mechanisms of Evasive Resistance in Cancer (MERiC), by The Swiss Initiative in Systems Biology grant SystemX – MERIC to M.H.H and by the Agence Nationale pour la Recherche (ANR JCJC 2017 LncEvoSys). C.K.Y.N. is supported by the Swiss Cancer League (KFS-4543-08-2018). This work was performed using the computing facilities of the CC LBBE/PRABI. We would also like to thank the French Institute of Bioinformatics (IFB, ANR-11-INBS-0013) for providing storage and computing resources on its national life science Cloud.

## Supplementary figure legends

**Supplementary Figure 1.**
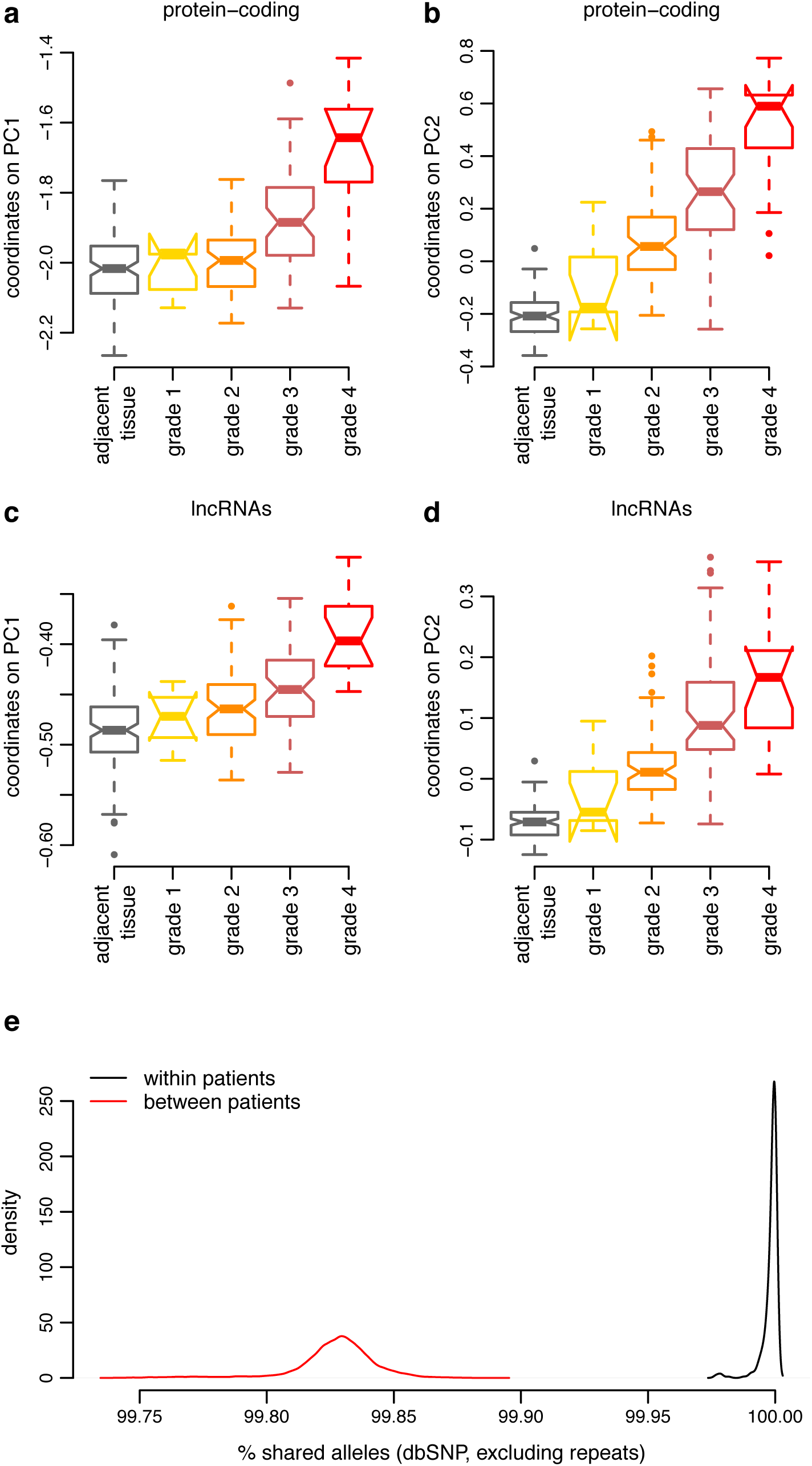
Sample clustering based on gene expression and genetic similarity. a. Boxplots representing the distribution of sample coordinates on principal component 1, for the PCA performed on protein-coding genes (displayed in figure 1). Samples are grouped depending on tissue types. Gray: adjacent tissue samples; yellow to red: tumors grouped by Edmondson-Steiner grade. Horizontal segments represent median values; notches represent 95% confidence intervals for the median; dashed segments extend to 1.5 times the inter-quartile range. b. Same as a, for principal component 2. c. Same as a, for the PCA performed on lncRNAs (displayed in figure 1). d. Same as c, for principal component 2. e. Distribution of the proportion of shared alleles for pairs of samples, for single nucleotide polymorphisms detected with our RNA-seq data (Methods). Red: distribution observed for pairs of samples derived from different patients; black: distribution observed from pairs of samples derived from the same patient.

**Supplementary Figure 2.**
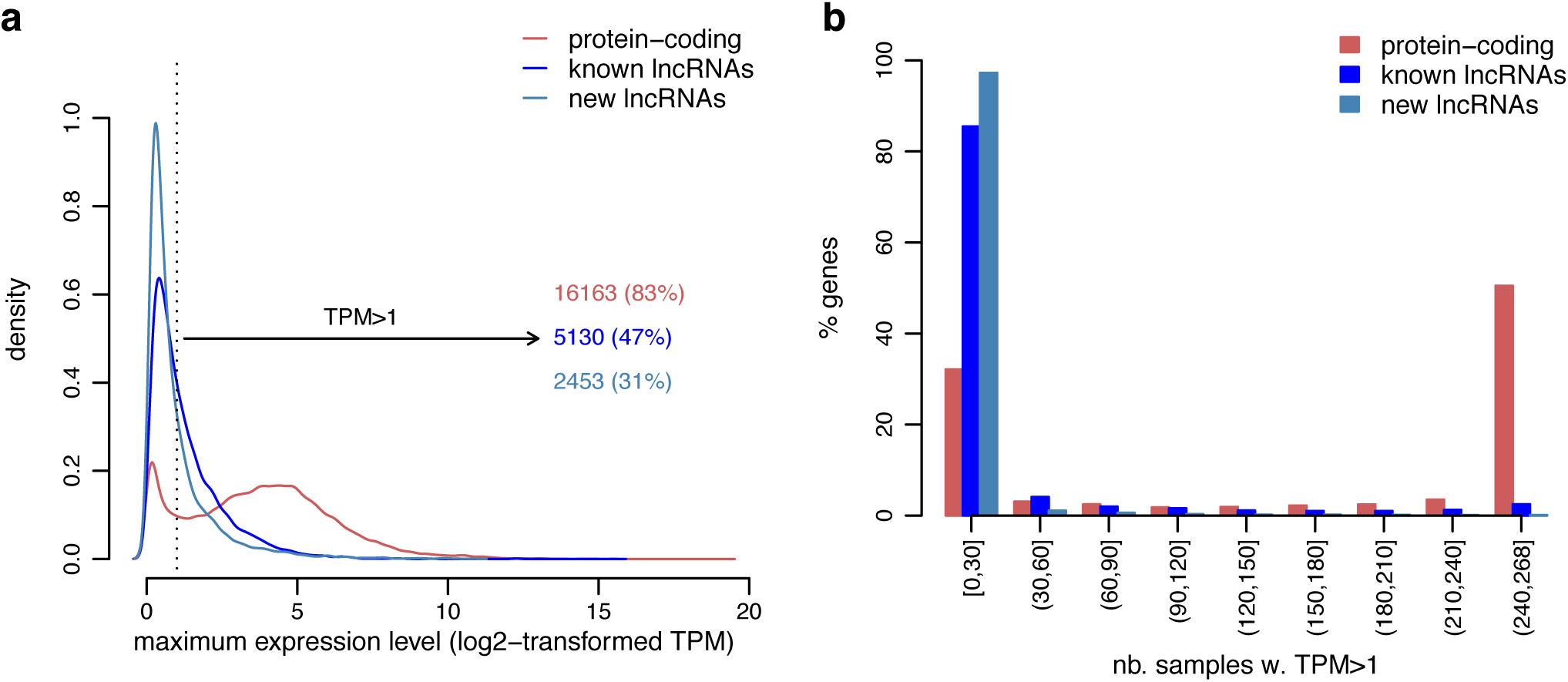
Expression patterns of protein-coding genes and lncRNAs in HCC samples. a. Distribution of the maximum expression level (log2-transformed TPM, maximum observed across samples) for protein-coding genes (red), previously known lncRNAs (dark blue) and newly annotated lncRNAs (light blue). The dotted vertical line represents the TPM = 1 threshold. Numbers of genes above the threshold are shown in the figure legend. b. Histogram of the number of samples in which the expression level is above the TPM = 1 threshold, for the three categories of genes described in a.

**Supplementary Figure 3.**
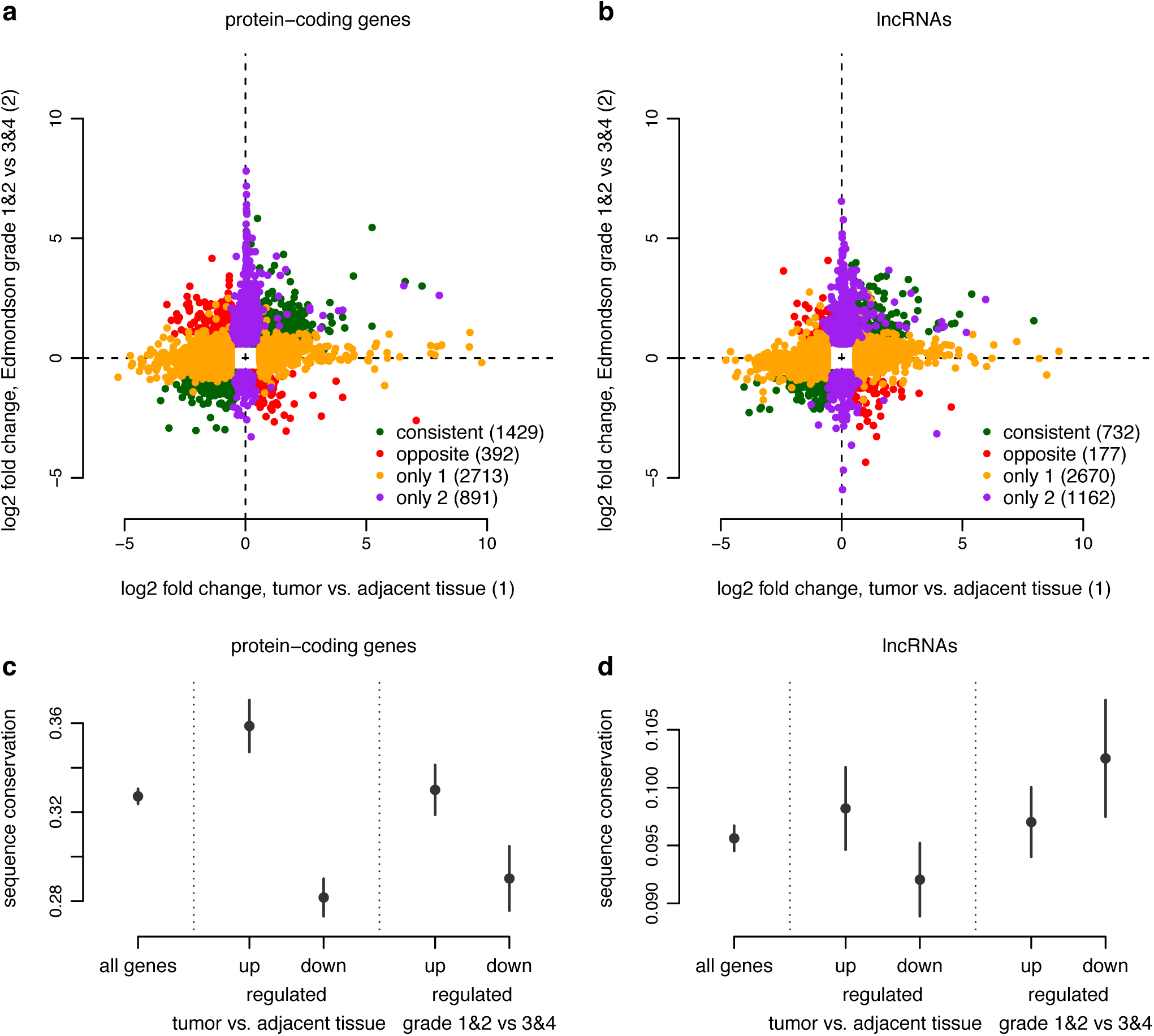
Differential expression patterns in HCC samples. a. Comparison between the log2 expression fold changes observed for our two main differential expression analyses (tumors vs. adjacent tissue samples, tumors with Edmondson-Steiner grades 3 and 4 vs. tumors with Edmondson-Steiner grades 1 and 2), for protein-coding genes. We show only genes that were significantly DE with a maximum FDR of 0.01 and a fold expression change above 1.5 in at least one of the two analyses. Green: genes with consistent expression changes in the both analyses; red: genes with opposite expression changes; orange: genes that are significantly DE only in the first DE analysis; purple: genes that are significantly DE only in the second DE analysis. b. Same as a, for lncRNAs. c. Distribution of sequence conservation scores for exonic regions (Methods), for protein-coding genes. Genes that are up-regulated or down-regulated in our two main DE analyses are shown separately. The dot represents the median conservation score, the vertical segments represents the 95% confidence interval for the median. d. Same as c, for lncRNAs.

**Supplementary Figure 4.**
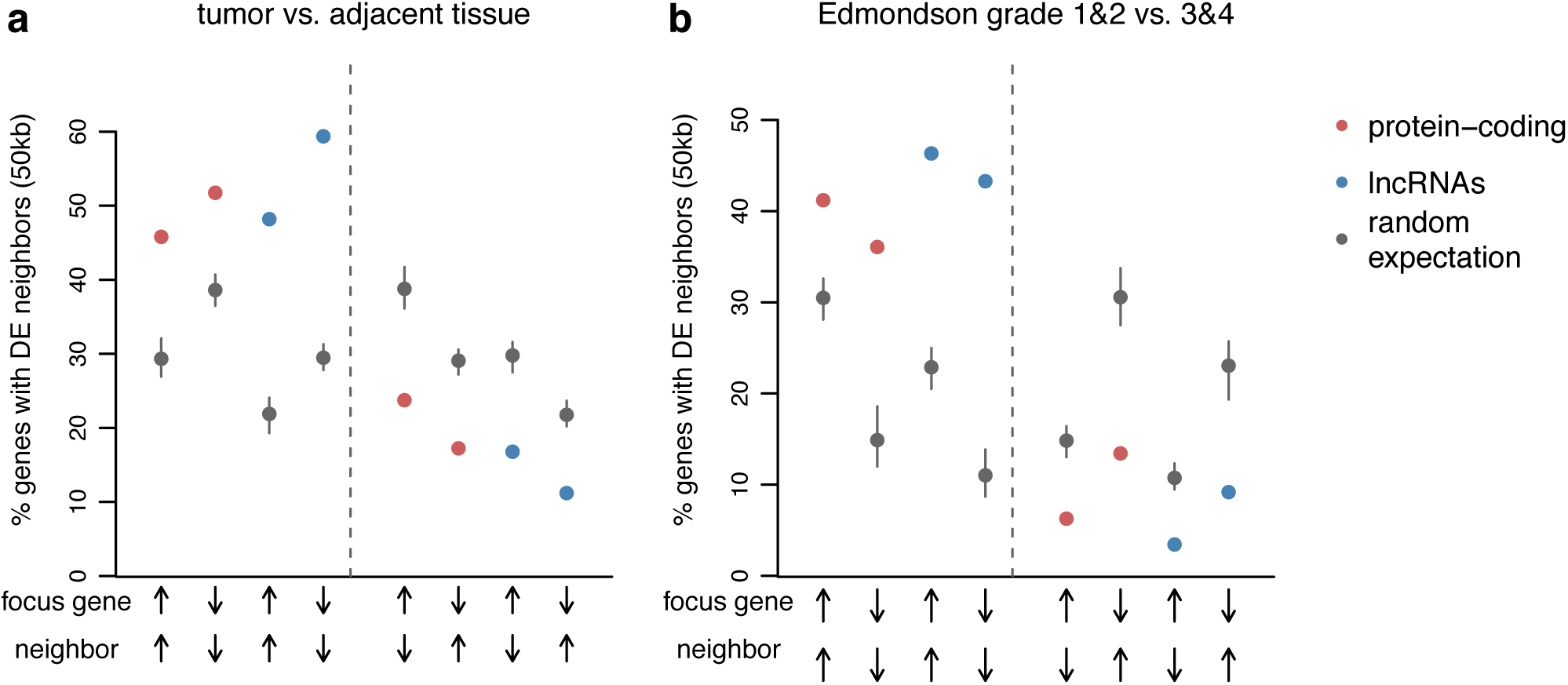
Genomic clustering of differentially expressed genes. a. Proportion of differentially expressed genes (maximum FDR 0.01, minimum fold expression change 1.5), in the comparison between paired tumors and adjacent samples, that have another differentially expressed gene within a 50kb distance. Red dots represent the values observed for protein-coding genes, blue dots for lncRNAs. The gray dots and vertical intervals represent the average and the 95% confidence intervals for the random expectation, obtained through simulations (Methods). The direction of the expression change required for the focus gene and the neighboring gene is displayed below the plot. b. Same as a, for the comparison between tumors with Edmondson-Steiner grades 3 and 4 vs. tumors with Edmondson-Steiner grades 1 and 2.

**Supplementary Figure 5.**
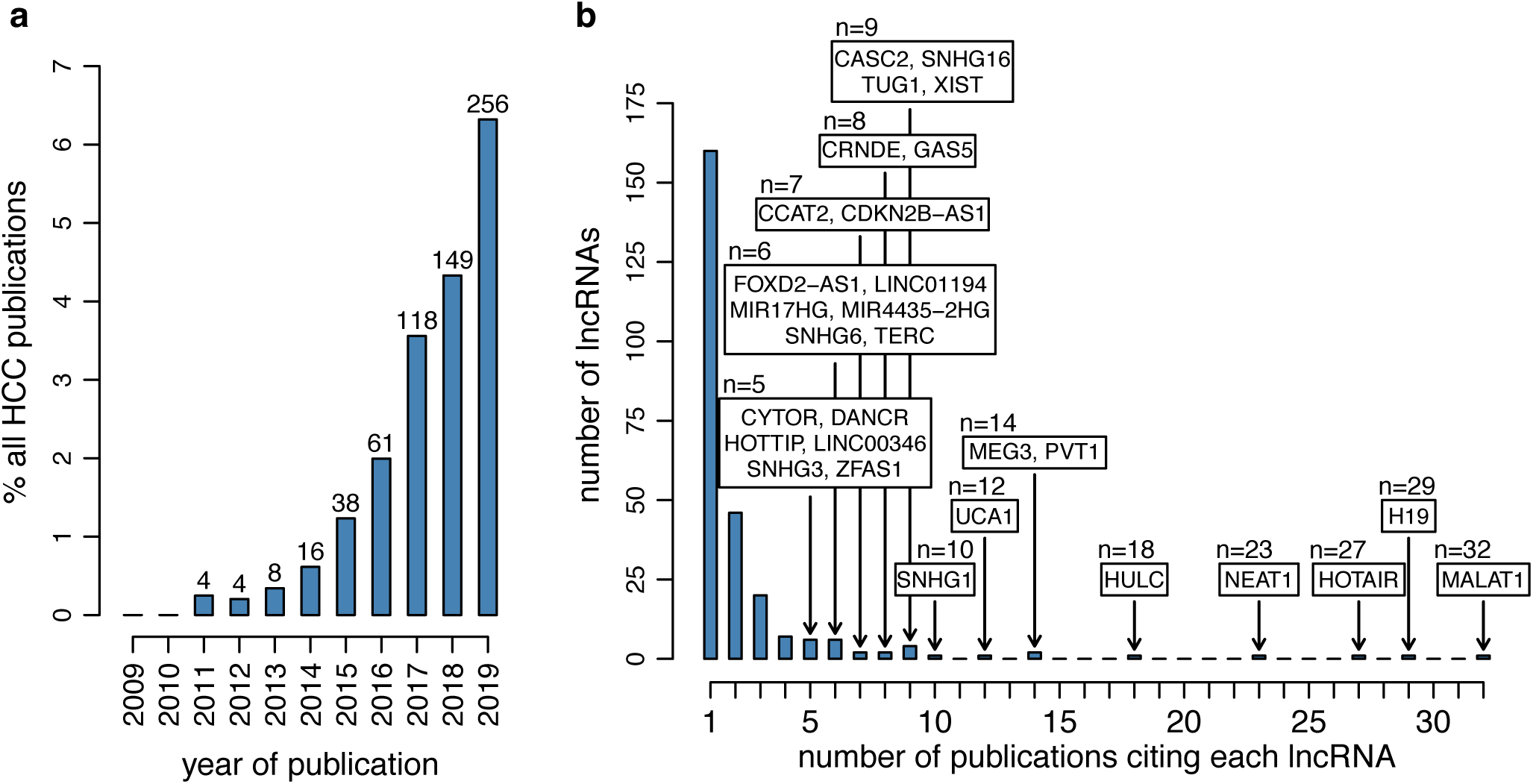
Growing interest for lncRNAs in the HCC field. a. Bar plot of the fraction of publications that mention lncRNAs and HCC, from 2009 to 2019. The bars represent the percentage of publications that mention lncRNAs, out of the total number of HCC publications. The numbers of publications that mention lncRNAs are shown above the bars. b. Histogram of the number of publications that cite each lncRNA in the context of HCC. lncRNAs that are cited in 5 or more publications are indicated in the plot.

**Supplementary Figure 6.**
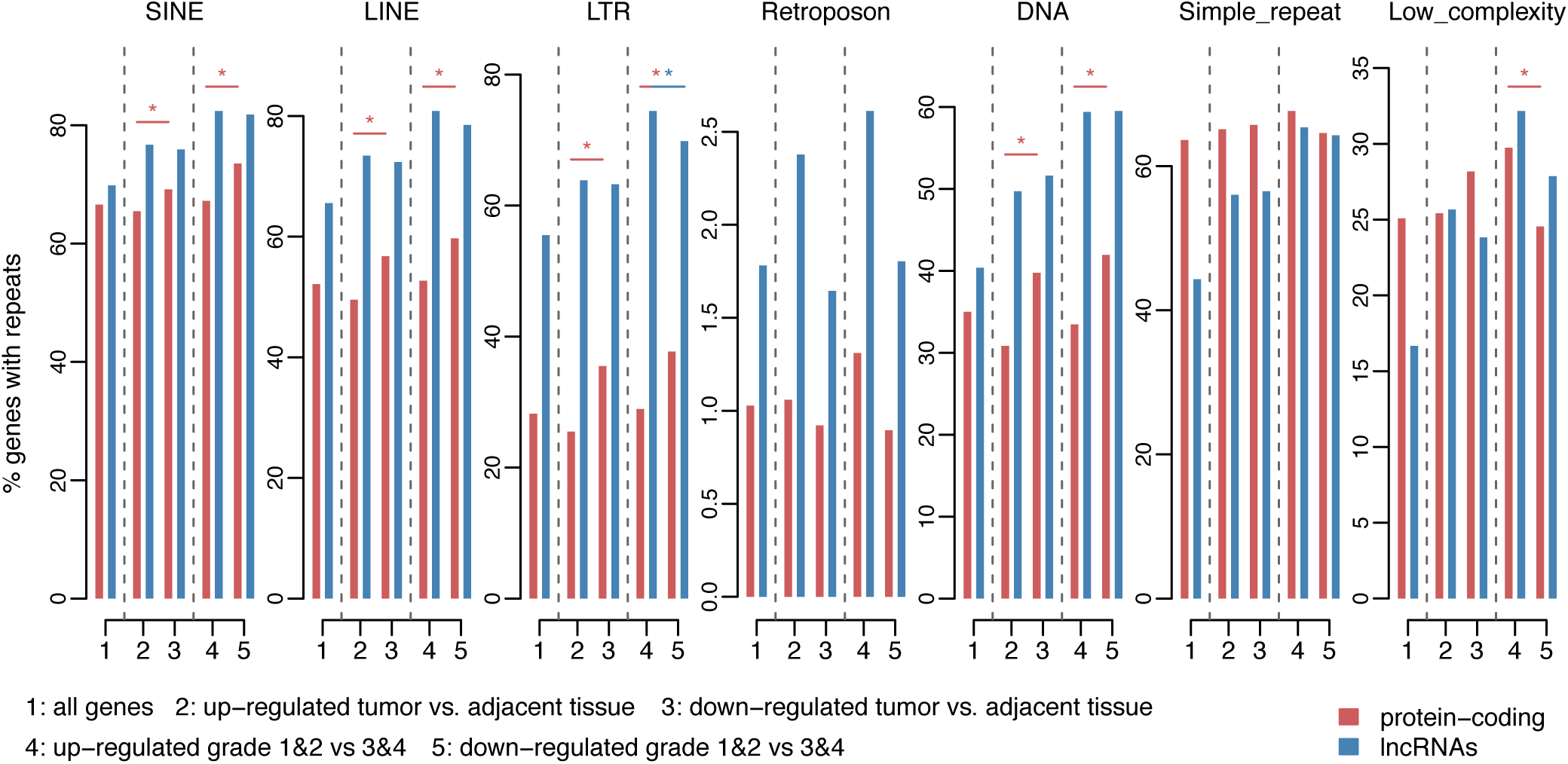
Increased repetitive sequence content in tumor-upregulated lncRNAs. Percentage of genes that have exonic overlap with major classes of repeats, for protein coding genes (red) and lncRNAs (blue). We display separately genes that show significant expression differences in our two main DE analyses. Significantly different proportions (Chi-square test, p-value <0.05) are marked by an asterisk.

**Supplementary Figure 7.**
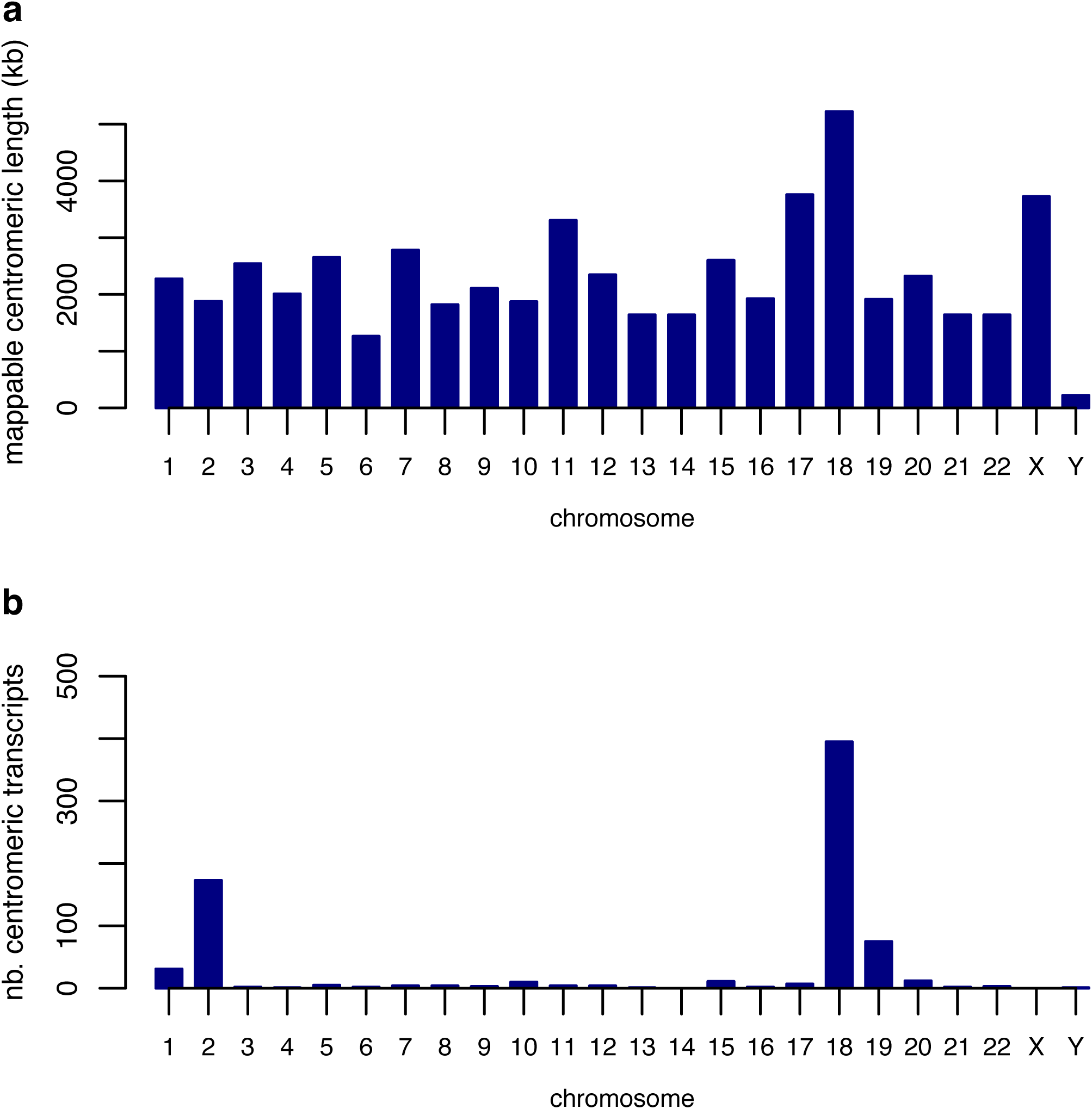
Centromeric transcription characteristics. a. Bar plot representing the total mappable length of centromeric regions, for each chromosome (Methods). b. Bar plot representing the number of transcribed loci found in centromeric regions, annotated with our RNA-seq data.

## Supplementary tables

Supplementary Table 1. Description of the 268 RNA-seq samples in our transcriptome collection.

Supplementary Table 2. List of cell type-specific markers for the most abundant cells in the healthy liver.

Supplementary Table 3. Results of our two main differential expression analyses for protein-coding genes and lncRNAs.

Supplementary Table 4. Gene ontology enrichment for differentially expressed protein-coding genes.

Supplementary Table 5. Number of HCC-related articles that mention each protein-coding and lncRNA genes.

Supplementary Table 6. Statistics for the overlap with different classes of repetitive elements.

Supplementary Table 7. Correlation between protein-coding gene expression and centromeric transcript levels across patients.

Supplementary Table 8. Results of our two main differential expression analyses for protein-coding genes involved in centromere functions.

## Supplementary datasets

Supplementary Dataset 1. Genome annotation used in this analysis, obtained by combining annotations from Ensembl 99 and gene models detected *de novo* with our RNA-seq data.

Supplementary Dataset 2. Gene expression data.

Supplementary Dataset 3. Full results of the differential expression analyses.

Supplementary Dataset 4. Evolutionary sequence conservation and repetitive element overlap statistics.

Supplementary Dataset 5. Mappable region coordinates and expression estimates for centromeric regions.

## References

Amândio AR, Necsulea A, Joye E, Mascrez B & Duboule D (2016) Hotair is dispensible for mouse development. PLoS Genet 12: e1006232

Arun G, Diermeier S, Akerman M, Chang K-C, Wilkinson JE, Hearn S, Kim Y, MacLeod AR, Krainer AR, Norton L, et al (2016) Differentiation of mammary tumors and reduction in metastasis upon Malat1 lncRNA loss. Genes Dev 30: 34–51

Bartolomei MS, Zemel S & Tilghman SM (1991) Parental imprinting of the mouse H19 gene. Nature 351: 153–155

Bouzinba-Segard H, Guais A & Francastel C (2006) Accumulation of small murine minor satellite transcripts leads to impaired centromeric architecture and function. Proc Natl Acad Sci USA 103: 8709–8714

Boyault S, Rickman DS, de Reyniès A, Balabaud C, Rebouissou S, Jeannot E, Hérault A, Saric J, Belghiti J, Franco D, et al (2007) Transcriptome classification of HCC is related to gene alterations and to new therapeutic targets. Hepatology 45: 42–52

Brawand D, Soumillon M, Necsulea A, Julien P, Csárdi G, Harrigan P, Weier M, Liechti A, Aximu-Petri A, Kircher M, et al (2011) The evolution of gene expression levels in mammalian organs. Nature 478: 343–348

Bray NL, Pimentel H, Melsted P & Pachter L (2016) Near-optimal probabilistic RNA-seq quantification. Nat Biotechnol 34: 525–527

Buchfink B, Xie C & Huson DH (2015) Fast and sensitive protein alignment using DIAMOND. Nat Methods 12: 59–60

Burns KH (2017) Transposable elements in cancer. Nature Reviews Cancer 17: 415–424

Bury L, Moodie B, Ly J, McKay LS, Miga KH & Cheeseman IM (2020) Alpha-satellite RNA transcripts are repressed by centromere-nucleolus associations. Elife 9

Carninci P, Kasukawa T, Katayama S, Gough J, Frith MC, Maeda N, Oyama R, Ravasi T, Lenhard B, Wells C, et al (2005) The transcriptional landscape of the mammalian genome. Science 309: 1559–1563

Casper J, Zweig AS, Villarreal C, Tyner C, Speir ML, Rosenbloom KR, Raney BJ, Lee CM, Lee BT, Karolchik D, et al (2018) The UCSC Genome Browser database: 2018 update. Nucleic Acids Res 46: D762–D769

Chen S, Wang G, Tao K, Cai K, Wu K, Ye L, Bai J, Yin Y, Wang J, Shuai X, et al (2020) Long noncoding RNA metastasis-associated lung adenocarcinoma transcript 1 cooperates with enhancer of zeste homolog 2 to promote hepatocellular carcinoma development by modulating the microRNA-22/Snail family transcriptional repressor 1 axis. Cancer Sci 111: 1582–1595

Chiatante G, Giannuzzi G, Calabrese FM, Eichler EE & Ventura M (2017) Centromere Destiny in Dicentric Chromosomes: New Insights from the Evolution of Human Chromosome 2 Ancestral Centromeric Region. Mol Biol Evol 34: 1669–1681

Cui H, Zhang Y, Zhang Q, Chen W, Zhao H & Liang J (2017) A comprehensive genome-wide analysis of long noncoding RNA expression profile in hepatocellular carcinoma. Cancer Med 6: 2932–2941

Cunningham F, Achuthan P, Akanni W, Allen J, Amode MR, Armean IM, Bennett R, Bhai J, Billis K, Boddu S, et al (2019) Ensembl 2019. Nucleic Acids Res 47: D745–D751

Darbellay F & Necsulea A (2020) Comparative transcriptomics analyses across species, organs, and developmental stages reveal functionally constrained lncRNAs. Mol Biol Evol 37: 240–259

Dray S & Dufour A (2007) The ade4 package: implementing the duality diagram for ecologists. Journal of Statistical Software 22: 1–20

Eden E, Navon R, Steinfeld I, Lipson D & Yakhini Z (2009) GOrilla: a tool for discovery and visualization of enriched GO terms in ranked gene lists. BMC Bioinformatics 10: 48

El-Gebali S, Mistry J, Bateman A, Eddy SR, Luciani A, Potter SC, Qureshi M, Richardson LJ, Salazar GA, Smart A, et al (2019) The Pfam protein families database in 2019. Nucleic Acids Res 47: D427–D432

Engreitz JM, Ollikainen N & Guttman M (2016) Long non-coding RNAs: spatial amplifiers that control nuclear structure and gene expression. Nat Rev Mol Cell Biol 17: 756–770

Ferri F, Bouzinba-Segard H, Velasco G, Hubé F & Francastel C (2009) Non-coding murine centromeric transcripts associate with and potentiate Aurora B kinase. Nucleic Acids Res 37: 5071–5080

Finn RS, Qin S, Ikeda M, Galle PR, Ducreux M, Kim T-Y, Kudo M, Breder V, Merle P, Kaseb AO, et al (2020) Atezolizumab plus bevacizumab in unresectable hepatocellular carcinoma. N Engl J Med 382: 1894–1905

Gupta RA, Shah N, Wang KC, Kim J, Horlings HM, Wong DJ, Tsai M-C, Hung T, Argani P, Rinn JL, et al (2010) Long non-coding RNA HOTAIR reprograms chromatin state to promote cancer metastasis. Nature 464: 1071–1076

Gutschner T & Diederichs S (2012) The hallmarks of cancer: a long non-coding RNA point of view. RNA Biol 9: 703–719

Hanahan D & Weinberg RA (2011) Hallmarks of cancer: the next generation. Cell 144: 646–674

Hao Y, Crenshaw T, Moulton T, Newcomb E & Tycko B (1993) Tumour-suppressor activity of H19 RNA. Nature 365: 764–767

Hartke J, Johnson M & Ghabril M (2017) The diagnosis and treatment of hepatocellular carcinoma. Semin Diagn Pathol 34: 153–159

Hartley G & O’Neill RJ (2019) Centromere repeats: hidden gems of the genome. Genes (Basel) 10

Hori T, Cao J, Nishimura K, Ariyoshi M, Arimura Y, Kurumizaka H & Fukagawa T (2020) Essentiality of CENP-A depends on its binding mode to HJURP. Cell Rep 33: 108388

Hoshida Y, Nijman SMB, Kobayashi M, Chan JA, Brunet J-P, Chiang DY, Villanueva A, Newell P, Ikeda K, Hashimoto M, et al (2009) Integrative transcriptome analysis reveals common molecular subclasses of human hepatocellular carcinoma. Cancer Res 69: 7385–7392

Hou Z, Xu X, Zhou L, Fu X, Tao S, Zhou J, Tan D & Liu S (2017) The long non-coding RNA MALAT1 promotes the migration and invasion of hepatocellular carcinoma by sponging miR-204 and releasing SIRT1. Tumour Biol 39: 1010428317718135

Hutchinson JN, Ensminger AW, Clemson CM, Lynch CR, Lawrence JB & Chess A (2007) A screen for nuclear transcripts identifies two linked noncoding RNAs associated with SC35 splicing domains. BMC Genomics 8: 39

Iyer MK, Niknafs YS, Malik R, Singhal U, Sahu A, Hosono Y, Barrette TR, Prensner JR, Evans JR, Zhao S, et al (2015) The landscape of long noncoding RNAs in the human transcriptome. Nat Genet 47: 199–208

Ji P, Diederichs S, Wang W, Böing S, Metzger R, Schneider PM, Tidow N, Brandt B, Buerger H, Bulk E, et al (2003) MALAT-1, a novel noncoding RNA, and thymosin beta4 predict metastasis and survival in early-stage non-small cell lung cancer. Oncogene 22: 8031–8041

Jin Y, Lee WY, Toh ST, Tennakoon C, Toh HC, Chow PK-H, Chung AY-F, Chong SS, Ooi LL-P-J, Sung W-K, et al (2019) Comprehensive analysis of transcriptome profiles in hepatocellular carcinoma. J Transl Med 17: 273

Keniry A, Oxley D, Monnier P, Kyba M, Dandolo L, Smits G & Reik W (2012) The H19 lincRNA is a developmental reservoir of miR-675 that suppresses growth and Igf1r. Nat Cell Biol 14: 659–665

Kim D, Langmead B & Salzberg SL (2015) HISAT: a fast spliced aligner with low memory requirements. Nat Methods 12: 357–360

Kim J, Piao H-L, Kim B-J, Yao F, Han Z, Wang Y, Xiao Z, Siverly AN, Lawhon SE, Ton BN, et al (2018) Long noncoding RNA MALAT1 suppresses breast cancer metastasis. Nat Genet 50: 1705–1715

Kishikawa T, Otsuka M, Suzuki T, Seimiya T, Sekiba K, Ishibashi R, Tanaka E, Ohno M, Yamagami M & Koike K (2018) Satellite RNA increases DNA damage and accelerates tumor formation in mouse models of pancreatic cancer. Mol Cancer Res 16: 1255–1262

Kishikawa T, Otsuka M, Yoshikawa T, Ohno M, Ijichi H & Koike K (2016) Satellite RNAs promote pancreatic oncogenic processes via the dysfunction of YBX1. Nat Commun 7: 13006

Kornienko AE, Dotter CP, Guenzl PM, Gisslinger H, Gisslinger B, Cleary C, Kralovics R, Pauler FM & Barlow DP (2016) Long non-coding RNAs display higher natural expression variation than protein-coding genes in healthy humans. Genome Biol 17: 14

Kou J-T, Ma J, Zhu J-Q, Xu W-L, Liu Z, Zhang X-X, Xu J-M, Li H, Li X-L & He Q (2020) LncRNA NEAT1 regulates proliferation, apoptosis and invasion of liver cancer. Eur Rev Med Pharmacol Sci 24: 4152–4160

Lai M, Yang Z, Zhou L, Zhu Q, Xie H, Zhang F, Wu L, Chen L & Zheng S (2012) Long non-coding RNA MALAT-1 overexpression predicts tumor recurrence of hepatocellular carcinoma after liver transplantation. Med Oncol 29: 1810–1816

Lander ES, Linton LM, Birren B, Nusbaum C, Zody MC, Baldwin J, Devon K, Dewar K, Doyle M, FitzHugh W, et al (2001) Initial sequencing and analysis of the human genome. Nature 409: 860–921

Lanzafame M, Bianco G, Terracciano LM, Ng CKY & Piscuoglio S (2018) The role of long non-coding RNAs in hepatocarcinogenesis. Int J Mol Sci 19

Li G, Shi H, Wang X, Wang B, Qu Q, Geng H & Sun H (2019) Identification of diagnostic long non-coding RNA biomarkers in patients with hepatocellular carcinoma. Mol Med Rep 20: 1121–1130

Li Z, Liu B, Jin W, Wu X, Zhou M, Liu VZ, Goel A, Shen Z, Zheng L & Shen B (2018) hDNA2 nuclease/helicase promotes centromeric DNA replication and genome stability. EMBO J 37

Liao Y, Smyth GK & Shi W (2019) The R package Rsubread is easier, faster, cheaper and better for alignment and quantification of RNA sequencing reads. Nucleic Acids Res

Lin R, Maeda S, Liu C, Karin M & Edgington TS (2007) A large noncoding RNA is a marker for murine hepatocellular carcinomas and a spectrum of human carcinomas. Oncogene 26: 851–858

Liu S, Qiu J, He G, Liang Y, Wang L, Liu C & Pan H (2019) LncRNA MALAT1 acts as a miR-125a-3p sponge to regulate FOXM1 expression and promote hepatocellular carcinoma progression. J Cancer 10: 6649–6659

MacParland SA, Liu JC, Ma X-Z, Innes BT, Bartczak AM, Gage BK, Manuel J, Khuu N, Echeverri J, Linares I, et al (2018) Single cell RNA sequencing of human liver reveals distinct intrahepatic macrophage populations. Nat Commun 9: 4383

Matouk IJ, DeGroot N, Mezan S, Ayesh S, Abu-lail R, Hochberg A & Galun E (2007) The H19 non-coding RNA is essential for human tumor growth. PLoS ONE 2: e845

Mattick JS & Makunin IV (2006) Non-coding RNA. Hum Mol Genet 15 Spec No 1: R17–29

Melé M, Mattioli K, Mallard W, Shechner DM, Gerhardinger C & Rinn JL (2017) Chromatin environment, transcriptional regulation, and splicing distinguish lincRNAs and mRNAs. Genome Res 27: 27–37

Mudge JM, Jungreis I, Hunt T, Gonzalez JM, Wright JC, Kay M, Davidson C, Fitzgerald S, Seal R, Tweedie S, et al (2019) Discovery of high-confidence human protein-coding genes and exons by whole-genome PhyloCSF helps elucidate 118 GWAS loci. Genome Res

Necsulea A, Soumillon M, Warnefors M, Liechti A, Daish T, Grutzner F & Kaessmann H (2014) The evolution of lncRNA repertoires and expression patterns in tetrapods. Nature 505: 635–640

Nojima T, Tellier M, Foxwell J, Ribeiro de Almeida C, Tan-Wong SM, Dhir S, Dujardin G, Dhir A, Murphy S & Proudfoot NJ (2018) Deregulated expression of mammalian lncRNA through loss of SPT6 induces R-loop formation, replication stress, and cellular senescence. Mol Cell 72: 970–984.e7

Panzitt K, Tschernatsch MMO, Guelly C, Moustafa T, Stradner M, Strohmaier HM, Buck CR, Denk H, Schroeder R, Trauner M, et al (2007) Characterization of HULC, a novel gene with striking up-regulation in hepatocellular carcinoma, as noncoding RNA. Gastroenterology 132: 330–342

Pertea M, Pertea GM, Antonescu CM, Chang T-C, Mendell JT & Salzberg SL (2015) StringTie enables improved reconstruction of a transcriptome from RNA-seq reads. Nat Biotechnol 33: 290–295

Pertea M, Shumate A, Pertea G, Varabyou A, Breitwieser FP, Chang Y-C, Madugundu AK, Pandey A & Salzberg SL (2018) CHESS: a new human gene catalog curated from thousands of large-scale RNA sequencing experiments reveals extensive transcriptional noise. Genome Biol 19: 208

Pradeepa MM, McKenna F, Taylor GCA, Bengani H, Grimes GR, Wood AJ, Bhatia S & Bickmore WA (2017) Psip1/p52 regulates posterior Hoxa genes through activation of lncRNA Hottip. PLoS Genet 13: e1006677

Quagliata L, Matter MS, Piscuoglio S, Arabi L, Ruiz C, Procino A, Kovac M, Moretti F, Makowska Z, Boldanova T, et al (2014) Long noncoding RNA HOTTIP/HOXA13 expression is associated with disease progression and predicts outcome in hepatocellular carcinoma patients. Hepatology 59: 911–923

Quagliata L, Quintavalle C, Lanzafame M, Matter MS, Novello C, di Tommaso L, Pressiani T, Rimassa L, Tornillo L, Roncalli M, et al (2018) High expression of HOXA13 correlates with poorly differentiated hepatocellular carcinomas and modulates sorafenib response in in vitro models. Lab Invest 98: 95–105

Rinn JL, Kertesz M, Wang JK, Squazzo SL, Xu X, Brugmann SA, Goodnough LH, Helms JA, Farnham PJ, Segal E, et al (2007) Functional demarcation of active and silent chromatin domains in human HOX loci by noncoding RNAs. Cell 129: 1311–1323

Schlackow M, Nojima T, Gomes T, Dhir A, Carmo-Fonseca M & Proudfoot NJ (2017) Distinctive patterns of transcription and RNA processing for human lincRNAs. Mol Cell 65: 25–38

Schultheiss CS, Laggai S, Czepukojc B, Hussein UK, List M, Barghash A, Tierling S, Hosseini K, Golob-Schwarzl N, Pokorny J, et al (2017) The long non-coding RNA H19 suppresses carcinogenesis and chemoresistance in hepatocellular carcinoma. Cell Stress 1: 37–54

Sherry ST, Ward MH, Kholodov M, Baker J, Phan L, Smigielski EM & Sirotkin K (2001) dbSNP: the NCBI database of genetic variation. Nucleic Acids Res 29: 308–311

Siepel A, Bejerano G, Pedersen JS, Hinrichs AS, Hou M, Rosenbloom K, Clawson H, Spieth J, Hillier LW, Richards S, et al (2005) Evolutionarily conserved elements in vertebrate, insect, worm, and yeast genomes. Genome Res 15: 1034–50

Smit AFA, Hubley R & Green P (2003) RepeatMasker Open-4.0.

Soneson C, Love MI & Robinson MD (2015) Differential analyses for RNA-seq: transcript-level estimates improve gene-level inferences. F1000Res 4: 1521

Soumillon M, Necsulea A, Weier M, Brawand D, Zhang X, Gu H, Barthès P, Kokkinaki M, Nef S, Gnirke A, et al (2013) Cellular source and mechanisms of high transcriptome complexity in the mammalian testis. Cell Rep 3: 2179–2190

Spielmann M, Lupiáñez DG & Mundlos S (2018) Structural variation in the 3D genome. Nat Rev Genet 19: 453–467

Talbert PB & Henikoff S (2018) Transcribing centromeres: noncoding RNAs and kinetochore assembly. Trends Genet 34: 587–599

Tietze L & Kessler SM (2020) The good, the bad, the question-H19 in hepatocellular carcinoma. Cancers (Basel) 12

Ting DT, Lipson D, Paul S, Brannigan BW, Akhavanfard S, Coffman EJ, Contino G, Deshpande V, Iafrate AJ, Letovsky S, et al (2011) Aberrant overexpression of satellite repeats in pancreatic and other epithelial cancers. Science 331: 593–596

Topp CN, Zhong CX & Dawe RK (2004) Centromere-encoded RNAs are integral components of the maize kinetochore. Proc Natl Acad Sci USA 101: 15986–15991

Unfried JP, Serrano G, Suárez B, Sangro P, Ferretti V, Prior C, Boix L, Bruix J, Sangro B, Segura V, et al (2019) Identification of coding and long non-coding RNAs differentially expressed in tumors and preferentially expressed in healthy tissues. Cancer Res

UniProt Consortium (2019) UniProt: a worldwide hub of protein knowledge. Nucleic Acids Res 47: D506–D515

Van der Auwera GA, Carneiro MO, Hartl C, Poplin R, Del Angel G, Levy-Moonshine A, Jordan T, Shakir K, Roazen D, Thibault J, et al (2013) From FastQ data to high confidence variant calls: the Genome Analysis Toolkit best practices pipeline. Curr Protoc Bioinformatics 43: 11.10.1-11.10.33

Wang F, Ying H-Q, He B-S, Pan Y-Q, Deng Q-W, Sun H-L, Chen J, Liu X & Wang S-K (2015) Upregulated lncRNA-UCA1 contributes to progression of hepatocellular carcinoma through inhibition of miR-216b and activation of FGFR1/ERK signaling pathway. Oncotarget 6: 7899–7917

Washietl S, Kellis M & Garber M (2014) Evolutionary dynamics and tissue specificity of human long noncoding RNAs in six mammals. Genome Res 24: 616–28

Yan X, Hu Z, Feng Y, Hu X, Yuan J, Zhao SD, Zhang Y, Yang L, Shan W, He Q, et al (2015) Comprehensive genomic characterization of long non-coding RNAs across human cancers. Cancer Cell 28: 529–540

Yang JD & Roberts LR (2010) Hepatocellular carcinoma: A global view. Nat Rev Gastroenterol Hepatol 7: 448–458

Yang Y, Chen L, Gu J, Zhang H, Yuan J, Lian Q, Lv G, Wang S, Wu Y, Yang Y-CT, et al (2017) Recurrently deregulated lncRNAs in hepatocellular carcinoma. Nature Communications 8: 14421

Yoshimizu T, Miroglio A, Ripoche M-A, Gabory A, Vernucci M, Riccio A, Colnot S, Godard C, Terris B, Jammes H, et al (2008) The H19 locus acts in vivo as a tumor suppressor. Proc Natl Acad Sci USA 105: 12417–12422

Zhou Y, Fan R-G, Qin C-L, Jia J, Wu X-D & Zha W-Z (2019) LncRNA-H19 activates CDC42/PAK1 pathway to promote cell proliferation, migration and invasion by targeting miR-15b in hepatocellular carcinoma. Genomics 111: 1862–1872

Zhu A, Ibrahim JG & Love MI (2019) Heavy-tailed prior distributions for sequence count data: removing the noise and preserving large differences. Bioinformatics 35: 2084–2092

